# SMN controls neuromuscular junction integrity through U7 snRNP

**DOI:** 10.1101/2021.08.31.458410

**Authors:** Sarah Tisdale, Meaghan Van Alstyne, Christian M. Simon, George Z. Mentis, Livio Pellizzoni

## Abstract

The neuromuscular junction (NMJ) is an essential synapse for animal survival whose loss is a key hallmark of neurodegenerative diseases such as amyotrophic lateral sclerosis (ALS) and spinal muscular atrophy (SMA). While insights into the function of the causative genes implicate RNA dysregulation in NMJ pathogenesis, the RNA-mediated mechanisms controlling the biology of this specialized synapse that go awry in disease remain elusive. Here, we show that activity of the SMA-determining SMN protein in the assembly of U7 small nuclear ribonucleoprotein (snRNP), which functions in the 3**’**-end processing of replication-dependent histone mRNAs, is required for NMJ integrity. AAV9-mediated gene delivery of U7-specific Lsm10 and Lsm11 proteins selectively enhances U7 snRNP assembly, corrects histone mRNA processing defects, and rescues key structural and functional abnormalities of neuromuscular pathology in SMA mice - including NMJ denervation, reduced synaptic transmission, and skeletal muscle atrophy. Furthermore, U7 snRNP dysfunction induced by SMN deficiency drives selective loss of the synaptic organizing protein Agrin at NMJs innervating vulnerable axial muscles of SMA mice, revealing an unanticipated link between U7-dependent histone mRNA processing and motor neuron-derived expression of an essential factor for NMJ biology. Together, these findings establish a direct contribution of U7 snRNP dysfunction to the neuromuscular phenotype in SMA and the requirement of RNA-mediated histone gene regulation for maintaining functional synaptic connections between motor neurons and muscles.

---

The neuromuscular junction (NMJ) is a specialized chemical synapse that controls motor behaviors indispensable for human life such as breathing, swallowing, and locomotion. NMJ innervation and function are strongly affected in some of the most severe and often-fatal neuromuscular diseases such as spinal muscular atrophy (SMA)^1, 2^, amyotrophic lateral sclerosis (ALS)^3^, and myasthenia gravis^4^ as well as by exposure to lethal toxins and venoms^5^. Importantly, RNA dysregulation is emerging as a driver of neuromuscular pathology in ALS and SMA among other neurodegenerative diseases^2, 6^, but the precise mechanisms are poorly understood. Understanding this connection has the potential to uncover novel aspects of NMJ biology in health and disease that have fundamental biomedical relevance.

NMJ denervation of vulnerable axial and proximal muscles is a hallmark of SMA^1, 2^, which is caused by a ubiquitous deficiency in the survival motor neuron (SMN) protein due to homozygous deletion or mutation of the *SMN1* gene^7, 8^. SMN is part of a multi-functional protein complex with essential roles in RNA regulation^9,10^, including the assembly of small nuclear ribonucleoproteins (snRNPs) of the spliceosomes that catalyze U2- and U12-type pre-mRNA splicing^11, 12^ as well as U7 snRNP required for processing of histone mRNAs^13, 14^. However, which of the numerous functions of SMN account for NMJ denervation and muscle fiber atrophy remains to be established.

Histones are highly conserved nuclear proteins that facilitate chromatin compaction and are central to gene regulation. Metazoan replication-dependent histones are encoded by a unique class of genes whose mRNAs lack introns and poly(A) tails^15^. The only processing event to occur on histone precursor mRNA is a single 3’-end endonucleolytic cleavage that is both dependent on U7 snRNP function and required for proper regulation of histone gene expression^15, 16^. We previously demonstrated that SMN deficiency disrupts U7 snRNP biogenesis and histone mRNA 3’-end processing in SMA mice and human patients^14^, but a direct pathogenic role for perturbation of this RNA pathway in the disease process has not been established. Moreover, while it is widely recognized that histone mRNA 3’-end processing is key for dividing cells to ensure that proper amounts of histones are synthesized to match the requirement for genome replication during S phase^15^, this need is not shared by neurons because they are postmitotic and it is unknown whether U7 function is relevant for the biology of neurons.

Designed to address these outstanding questions through selective restoration of U7 snRNP biogenesis and function in a mouse model of SMA, this study identifies the specific SMN-dependent RNA pathway driving NMJ denervation in SMA and reveals an unexpected role for U7-mediated histone mRNA processing in motor neuron biology and disease.

## Lsm10 and Lsm11 co-expression enhances U7 snRNP assembly

SMN mediates the assembly of Sm proteins onto their respective snRNA substrates to generate functional snRNPs that carry out pre-mRNA splicing^11, 12^. In addition, specialized SMN complexes in which the Sm-like proteins Lsm10 and Lsm11 replace SmD1 and SmD2 of spliceosomal snRNPs mediate the assembly of U7 snRNP^13, 14, 17^ (Fig. 1a), which functions not in splicing but in histone mRNA processing ^15, 16^. These SMN complexes are limiting due to the low abundance of Lsm10 and Lsm11 proteins relative to the amounts of Sm proteins that are assembled on spliceosomal snRNPs^13, 17^. We therefore reasoned that increasing the levels of U7-specific Lsm10 and Lsm11 could be a means to selectively enhance U7 snRNP assembly. To test this, we generated NIH3T3-Smn_RNAi_ cell lines with doxycycline (Dox)-regulated RNAi knockdown of endogenous SMN^18, 19^ that overexpressed FLAG-tagged Lsm10 and Lsm11 either alone or in combination (Extended Data Fig. 1a-b). Immunoprecipitation experiments demonstrated that epitope-tagged Lsm10 and Lsm11 specifically incorporate into endogenous U7 snRNPs in these cell lines (Extended Data Fig. 1c). Using *in vitro* snRNP assembly assays with radioactive snRNAs^12, 14^, we found that co-expression of Lsm10 and Lsm11—but neither on its own—specifically and strongly increased U7 but not U1 snRNP assembly without changes in SMN levels (Fig. 1b, c). To determine the effects of Lsm10/11 overexpression *in vivo*, we first developed a single-cassette vector for co-expression of FLAG-tagged Lsm10 and Lsm11 that similarly enhanced U7 snRNP assembly *in vitro* (Extended Data Fig. 1d-f). We then delivered an adeno-associated virus serotype 9 (AAV9) vector harboring this cassette under a CAG promoter in the central nervous system (CNS) of WT mice at P0 by intracerebroventricular (ICV) injection followed by analysis of snRNP assembly at P11 (Extended Data Fig. 2a). AAV9-Lsm10/11 robustly increased U7 but not U1 snRNP assembly in brain extracts (Extended Data Fig. 2b-c). Thus, co-expression of Lsm10 and Lsm11 is necessary and sufficient to selectively increase SMN-dependent U7 snRNP assembly both *in vitro* and *in vivo*.

**Fig. 1.**
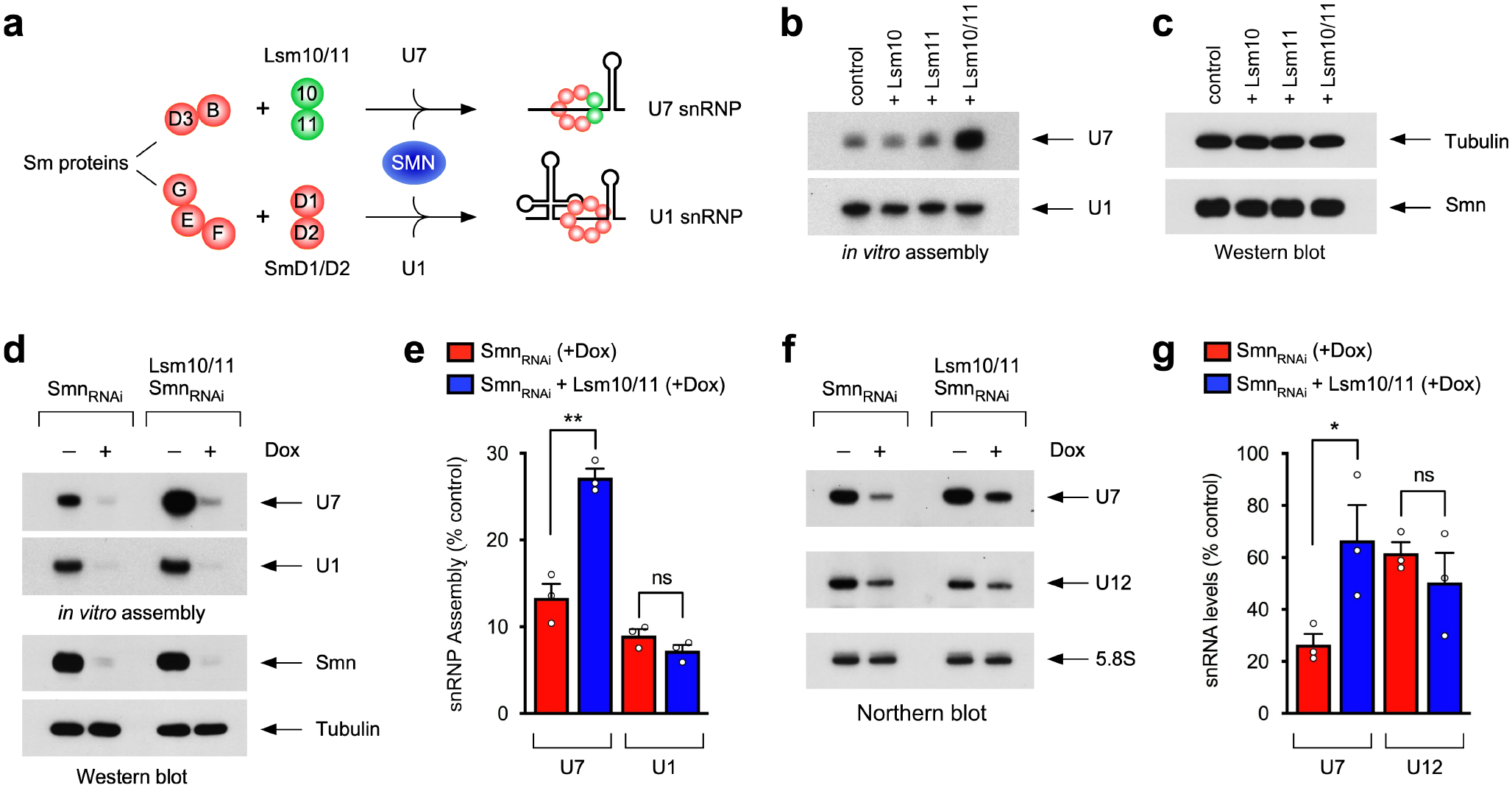
Co-expression of Lsm10 and Lsm11 promotes U7 snRNP assembly and prevents U7 snRNA reduction induced by SMN deficiency. **a**, Schematic representation of SMN-mediated assembly of U7 and U1 snRNAs with their respective Sm and Lsm proteins. **b**, U7 and U1 *in vitro* snRNP assembly with extracts from NIH3T3-Smn_RNAi_ cell lines with or without overexpression of FLAG-tagged Lsm10 and Lsm11 as indicated. **c**, Western blot analysis of SMN levels in cell extracts used in (**b**). **d**, U7 and U1 *in vitro* snRNP assembly (top panels) and Western blot analysis (bottom panels) with extracts from NIH3T3-Smn_RNAi_ and NIH3T3-Lsm10/11/Smn_RNAi_ cells treated with or without Dox for 5 days. **e**, Percentage of U7 and U1 snRNP assembly in Dox treated cells relative to control NIH3T3-Smn_RNAi_ cells without Dox from experiments as in (**d**). Data are mean and s.e.m (n=3 independent experiments). **P<0.01 (two-sided Student’s *t* test; P=0.002). NS, not significant (P>0.05) (two-sided Student’s *t* test; P=0.1573). **f**, Northern blot analysis of the indicated endogenous RNAs in the same experimental groups as in (**d**). **g**, Percentage of U7 and U1 snRNA levels in Dox treated cells relative to control NIH3T3-Smn_RNAi_ cells without Dox from experiments as in (**f**). Data are mean and s.e.m (n=3 independent experiments) normalized to 5.8S rRNA. *P<0.05 (two-sided Student’s *t* test; P=0.0472). NS, not significant (P>0.05) (two-sided Student’s *t* test; P=0.4069).

We next investigated the effects of Lsm10/11 co-expression when U7 biogenesis is strongly reduced by SMN deficiency^14^. Although both Smn_RNAi_ and Lsm10/11-Smn_RNAi_ cells display a similar Dox-dependent, RNAi-mediated knockdown of Smn to ∼10% of normal levels, Lsm10/11 co-expression led to a strong increase in the assembly of U7 but not U1 snRNP in SMN-deficient cells (Fig. 1d-e). Furthermore, Lsm10/11 co-expression specifically corrected the reduction in the endogenous levels of U7 but not U12 snRNA induced by SMN deficiency in NIH3T3 (Fig. 1f-g)^14, 18^. Therefore, Lsm10/11 co-expression can selectively increase both U7 snRNP assembly and steady-state U7 levels in SMN-deficient mammalian cells.

To determine the impact of Lsm10/11 co-expression on defective U7 biogenesis in a mouse model of SMA, we employed AAV9-mediated gene delivery of Lsm10/11 in SMN*Δ*7 mice^20^. While overexpression of Lsm10 and Lsm11 mRNAs did not alter the levels of SMN relative to GFP-treated SMA mice (Extended Data Fig. 2d-e), AAV9-Lsm10/11 partially corrected the severe reduction in U7 snRNP assembly induced by SMN deficiency (Extended Data Fig. 2f-g). Moreover, AAV9-Lsm10/11 significantly increased endogenous U7 snRNA levels in the liver of SMA mice (Extended Data Fig. 2h-i). Such an increase was below the threshold of detection in CNS tissue, likely reflecting the much more widespread transduction of liver than CNS by AAV9^21^. Thus, Lsm10/11 co-expression selectively enhances U7 snRNP biogenesis in SMA mice in the absence of SMN modulation.

## Selective correction of histone gene dysregulation by Lsm10/11 in SMA models

The only cellular function of U7 snRNP is its activity in the 3’-end processing of replication-dependent histone mRNAs^15, 16^, which is disrupted in SMA as a direct consequence of the reduced levels of U7 snRNP assembly induced by SMN deficiency^14^. Remarkably, Lsm10/11 co-expression robustly corrected the accumulation of uncleaved, 3’-extended histone mRNAs in SMN-deficient NIH3T3 cells (Fig. 2a) and in tissue of SMA mice (Fig. 2c and Extended Data Fig. 3d). The effects were specific because SMN-dependent RNA processing defects unrelated to U7 function^14, 18, 22^ were not improved (Extended Data Fig. 3). Thus, Lsm10/11 co-expression results in the selective correction of U7-mediated histone mRNA 3’-end processing deficits induced by SMN deficiency both *in vitro* and *in vivo*.

**Fig. 2.**
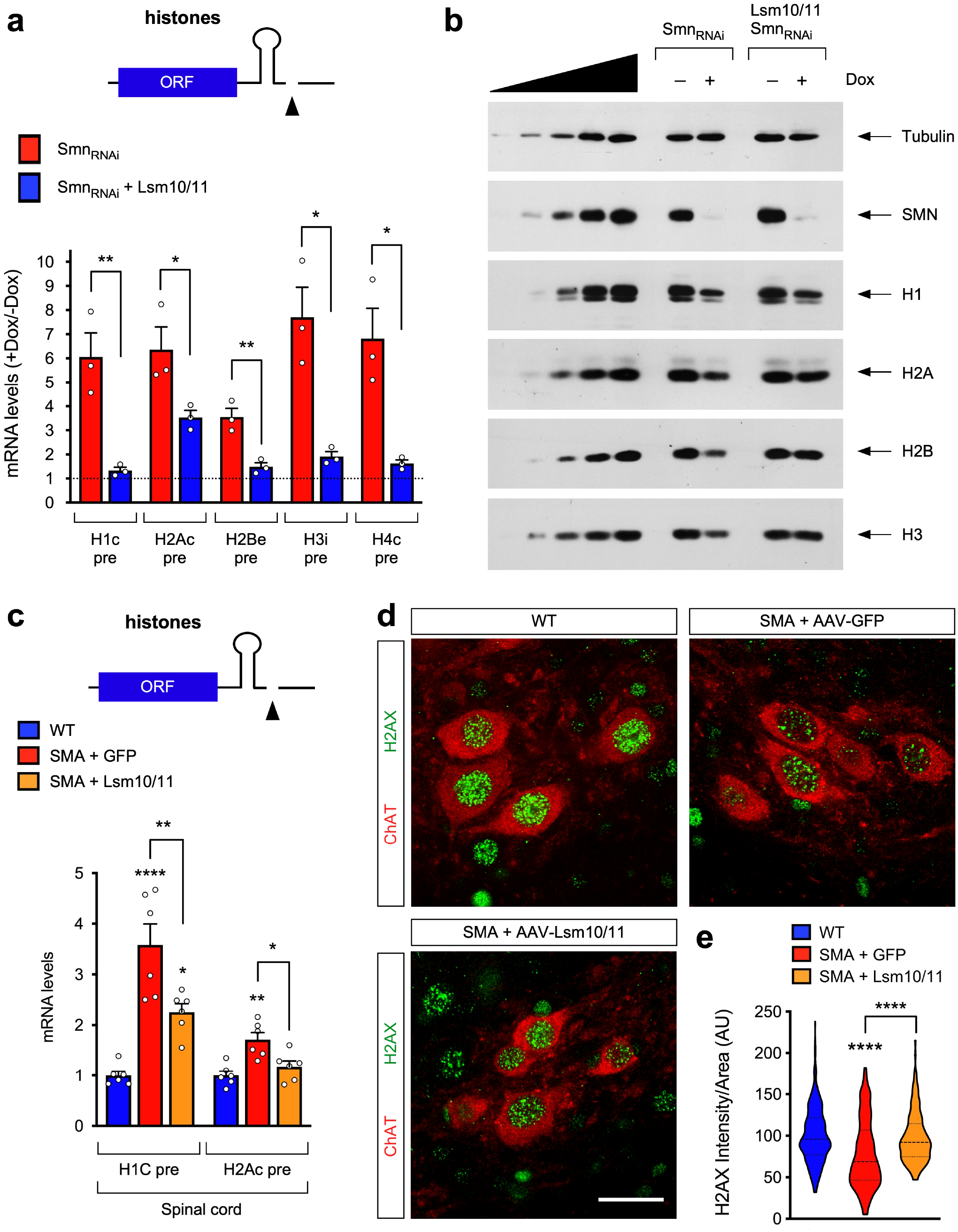
Co-expression of Lsm10 and Lsm11 corrects histone gene dysregulation induced by SMN deficiency. **a**, RT-qPCR analysis of 3’-extended histone mRNA levels in NIH3T3-Smn_RNAi_ and NIH3T3-Lsm10/11/Smn_RNAi_ cells. Schematic representation of a histone pre-mRNA and the 3’-end cleavage is shown at the top. For each cell line, fold change values in Dox treated cells relative to untreated cells set as 1 are shown. Data are mean and s.e.m (n=3 independent experiments) normalized to Gapdh mRNA. *P<0.05; **P<0.01 (two-way Student’s *t* test; P=0.0092 for H1c; P=0.0456 for H2Ac; P=0.0067 for H2Be; P=0.0101 for H3i; P=0.0148 for H4c). **b**, Western blot analysis of histone levels in NIH3T3-Smn_RNAi_ and NIH3T3-Lsm10/11/Smn_RNAi_ cells with or without Dox treatment. A two-fold serial dilution of control extract is shown on the left. **c**, RT-qPCR analysis of 3’-extended histone mRNAs in the spinal cord of SMA mice injected with AAV9-GFP or AAV9-Lsm10/11 relative to untreated WT mice at P11 set as 1. Schematic representation of a histone pre-mRNA and the 3’-end cleavage is shown at the top. Data are mean and s.e.m (n=6 mice) normalized to Gapdh mRNA. *P<0.05; **P<0.01; ***P<0.001 (one-way ANOVA with Tukey’s *post hoc* test; P<0.0001 for H1c in WT vs SMA+GFP; P=0.0109 for H1c in WT vs SMA+Lsm10/11; P=0.0073 for H1c in SMA+GFP vs SMA+Lsm10/11; P=0.0016 for H2Ac in WT vs SMA+GFP; P=0.0137 for H2Ac in SMA+GFP vs SMA+Lsm10/11). **d**, Histone H2AX and ChAT immunostaining of L1 spinal cords from untreated WT mice and SMA mice injected with AAV9-GFP or AAV9-Lsm10/11 at P11. Scale bar=20µm. **e**, Normalized fluorescent intensity of nuclear H2AX signal in ChAT^+^ L1 motor neurons from experiments as in (**d**). The violin plots show the median (solid line) and interquartile range (dotted lines) from the following number of motor neurons from 3 mice per group (n=295 neurons for WT; n=209 neurons for SMA+GFP; n=178 neurons SMA+Lsm10/11). ****P<0.0001 (one-way ANOVA with Tukey’s *post hoc* test; P<0.0001 for WT vs SMA+GFP; P<0.0001 for SMA+GFP vs SMA+Lsm10/11).

Lsm10/11 co-expression restored histone protein levels that are reduced in SMN-deficient NIH3T3 cells without an impact on cell proliferation (Fig. 2b and Extended Data Fig. 4a, b), indicating that they result directly from U7-dependent dysregulation of histone mRNA 3’-end processing and not indirectly from halted cell division. Given that histone levels were not globally affected in the spinal cord of WT and SMA mice at P11 (Extended Data Fig. 4c), we looked into disease-relevant motor neurons by immunostaining. Since each of the core histones represents a family of proteins encoded by multiple genes that cannot be distinguished using available antibodies^15^, we therefore performed immunostaining against H2AX – a single copy histone gene that undergoes U7-dependent 3’-end processing and is affected by SMN deficiency^14^. We found a reduction of H2AX expression in motor neurons from SMA mice injected with AAV9-GFP relative to WT motor neurons that was corrected by AAV9-Lsm10/11 injection (Fig. 2d, e). Thus, SMN deficiency can affect histone expression in SMA motor neurons and correction by Lsm10/11 indicates that this defect is downstream of U7 snRNP dysfunction.

## U7 snRNP dysfunction contributes to neuromuscular pathology in SMA mice

Having established Lsm10/11 co-expression as a means for the selective improvement of defective U7 biogenesis and histone mRNA 3’-end processing – which represent the most proximal molecular events linked to the dysfunction of SMN in this RNA pathway – we next sought to assess their contribution to the disease process. To do so, we investigated the effects of AAV9-Lsm10/11 injection on the phenotype of SMA mice using AAV9-GFP and AAV9-SMN as negative and positive controls, respectively. AAV9-Lsm10/11 moderately improved motor function in SMA mice without affecting either weight or survival compared to GFP-injected SMA mice (Extended Data Fig. 5a-c), while AAV9-SMN robustly corrected all parameters as expected^21, 23–25^.

To determine which elements of the severe neuromuscular phenotype of SMA mice could be improved by Lsm10/11, we first analyzed synaptic transmission at the NMJ by performing EMG recordings from the *quadratus lumborum* (QL) – a disease-relevant, axial muscle – upon evoked stimulation of the L1 ventral root using the *ex vivo* spinal cord preparation. Remarkably, the amplitude of the compound muscle action potential (CMAP) that is ablated in GFP-injected SMA mice relative to WT controls at P11 was strongly improved by AAV9-Lsm10/11 though not to the same extent as SMN restoration (Fig. 3a, b). The effects were specific to NMJ transmission because the reduction in the amplitude of the compound motor neuron output upon stimulation of proprioceptive afferent fibers in the L1 dorsal root was not improved by AAV9-Lsm10/11 in SMA mice (Extended Data Fig. 5d, e). Importantly, while about 40% of NMJs were fully denervated in the QL of SMA mice treated with AAV-GFP, this defect was robustly corrected by AAV9-Lsm10/11 (Fig. 3c, d). In contrast, the loss of VGluT1^+^ proprioceptive sensory synapses onto the somata of L1 motor neurons of GFP-treated SMA mice at P11 was rescued by AAV9-SMN but not AAV9-Lsm10/11 (Extended Data Fig. 6a and d). Lsm10/11 co-expression in SMA mice had also no effect on the loss of vulnerable L1 motor neurons (Extended Data Fig. 6b and e) and L5 medial motor column (MMC) motor neurons (Extended Data Fig. 6c and f). Altogether, these results demonstrate that enhancement of U7 snRNP biogenesis through Lsm10/11 co-expression strongly improves NMJ innervation and function in SMA mice, while the lack of correction of deafferentation and death of SMA motor neurons supports the selectivity of the effects because these pathogenic events have been linked to SMN-dependent splicing changes unrelated to U7 function^22, 25–27^.

**Fig. 3.**
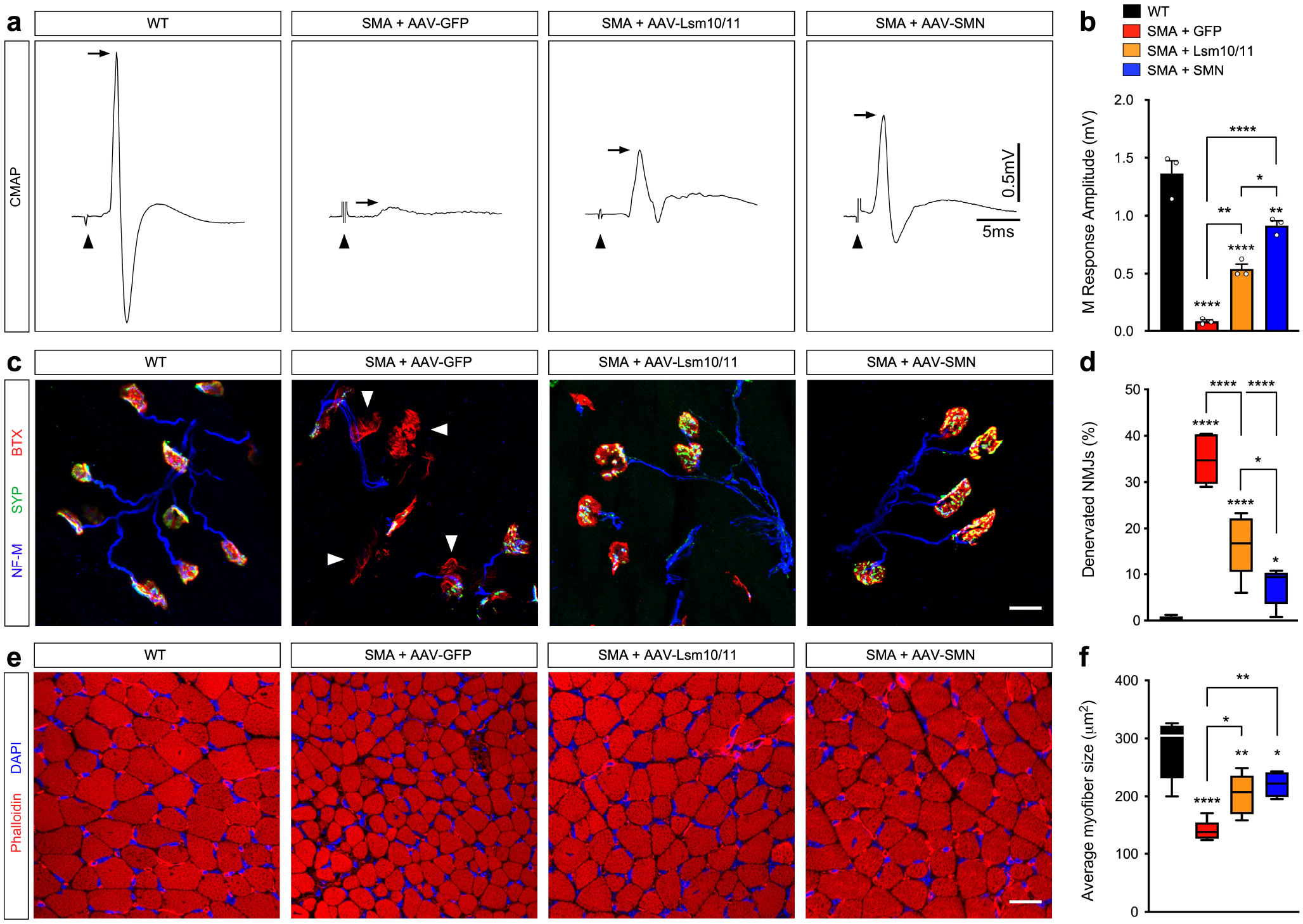
Co-expression of Lsm10 and Lsm11 corrects NMJ denervation and skeletal muscle atrophy in SMA mice. **a**, Representative EMG tracings of CMAP recorded from the *quadratus lumborum* muscle of untreated WT mice and SMA mice injected with AAV9-GFP, AAV9-Lsm10/11 or AAV9-SMN at P11. Arrows indicate peak CMAP amplitude and arrowheads indicate artifact from stimulus. **b**, Amplitude of the M-response from the same groups as in (**a**) at P11. Data are mean and s.e.m (n=3 mice). *P<0.05; **P<0.01; ****P<0.0001 (one-way ANOVA with Tukey’s *post hoc* test; P<0.0001 for WT vs SMA+GFP; P<0.0001 for WT vs SMA+Lsm10/11; P=0.0045 for WT vs SMA+SMN; P=0.0040 for SMA+GFP vs SMA+Lsm10/11; P<0.0001 for SMA+GFP vs SMA+SMN; P=0.0128 for SMA+Lsm10/11 vs SMA+SMN). **c**, NMJ immunostaining with Synaptophysin (SYP), Neurofilament-M (NF-M) and *α*-bungarotoxin (BTX) of *quadratus lumborum* muscles from the same groups as in (**a**) at P11. Denervated NMJs lacking pre-synaptic SYP and NF-M staining are indicated by arrowheads. Scale bar=20µm. **d**, Percentage of fully denervated NMJs from experiments as in (**c**). The box-and-whiskers graph shows the median, interquartile range, minimum and maximum from the following number of mice per group (n=7 for WT, and SMA+GFP; n=5 for SMA+Lsm10/11; n=6 for SMA+SMN). *P<0.05; ****P<0.0001 (one-way ANOVA with Tukey’s *post hoc* test; P<0.0001 for WT vs SMA+GFP; P<0.0001 for WT vs SMA+Lsm10/11; P=0.0337 for WT vs SMA+SMN; P<0.0001 for SMA+GFP vs SMA+Lsm10/11; P<0.0001 for SMA+GFP vs SMA+SMN; P=0.0119 for SMA+Lsm10/11 vs SMA+SMN). **e**, TRITC-conjugated phalloidin and DAPI staining of cross sections of the triceps muscle from the same groups as in (**a**) at P11. Scale bar=20µm. **f**, Quantification of the average myofiber size (μm^2^) from experiments as in (**e**). The box-and-whiskers graph shows the median, interquartile range, minimum and maximum from the following number of mice (n=5 for WT, SMA+Lsm10/11, and SMA+SMN; n=6 for SMA+GFP). *P<0.05; **P<0.01; ****P<0.0001 (one-way ANOVA with Tukey’s *post hoc* test; P<0.0001 for WT vs SMA+GFP; P=0.0093 for WT vs SMA+Lsm10/11; P=0.0463 for WT vs SMA+SMN; P=0.0319 for SMA+GFP vs SMA+Lsm10/11; P=0.0058 for SMA+GFP vs SMA+SMN).

Dysfunction and loss of NMJs contribute to skeletal muscle atrophy – a hallmark of SMA pathology^1, 2, 8^ – and the myofiber size in the *triceps brachii* is strongly reduced in AAV9-GFP-treated SMA mice relative to WT mice at P11 (Fig. 3e, f). Remarkably, treatment with AAV9-Lsm10/11, similar to AAV9-SMN, significantly improved this phenotype in SMA mice as further highlighted by an increased proportion of large-sized myofibers (Fig. 3e, f and Extended Data Fig. 7), revealing that U7 snRNP restoration improves skeletal muscle atrophy in SMA mice.

## SMN regulates Agrin expression at the NMJ through U7 snRNP

The identification of impaired U7 snRNP assembly as the upstream trigger of a cascade leading to motor neuron denervation and muscle atrophy in SMA mice raised the question as to which downstream effector(s) could directly mediate the distal effects of U7 dysfunction at the NMJ. A candidate emerging from recent studies^28–31^ was Agrin — an essential NMJ organizer that promotes clustering of post-synaptic acetylcholine receptors (AChR) during development^32, 33^ and is also required for postnatal NMJ maintenance^34^. Since increasing Agrin levels or its associated signaling cascade has been shown to ameliorate NMJ pathology of SMA mice in a manner similar to Lsm10/11 co-expression^28–31^, we sought to investigate possible links between Agrin and SMN-regulated U7 snRNP assembly. Motor neuron-derived Agrin isoforms (Z^+^ Agrin) have greatly increased potency in AChR clustering due to an amino acid insertion resulting from the inclusion of Z exons (Fig. 4a)^35, 36^. We found that, while Agrin mRNA levels were similar in the spinal cord of WT and SMA mice at P6, lumbar SMA motor neurons innervating the QL and iliopsoas muscles had reduced levels of both total and Z exon-skipped (*Δ*Z) Agrin mRNAs (Fig. 4a, b). The increased levels of 3’-extended H1c transcripts and Cdkn1a mRNA confirmed U7 snRNP dysfunction and RNA dysregulation in SMA motor neurons (Fig. 4b)^14, 19^. These results suggest that transcript level changes rather than previously reported splicing dysregulation^37^ may affect Agrin expression in vulnerable SMA motor neurons.

**Fig. 4.**
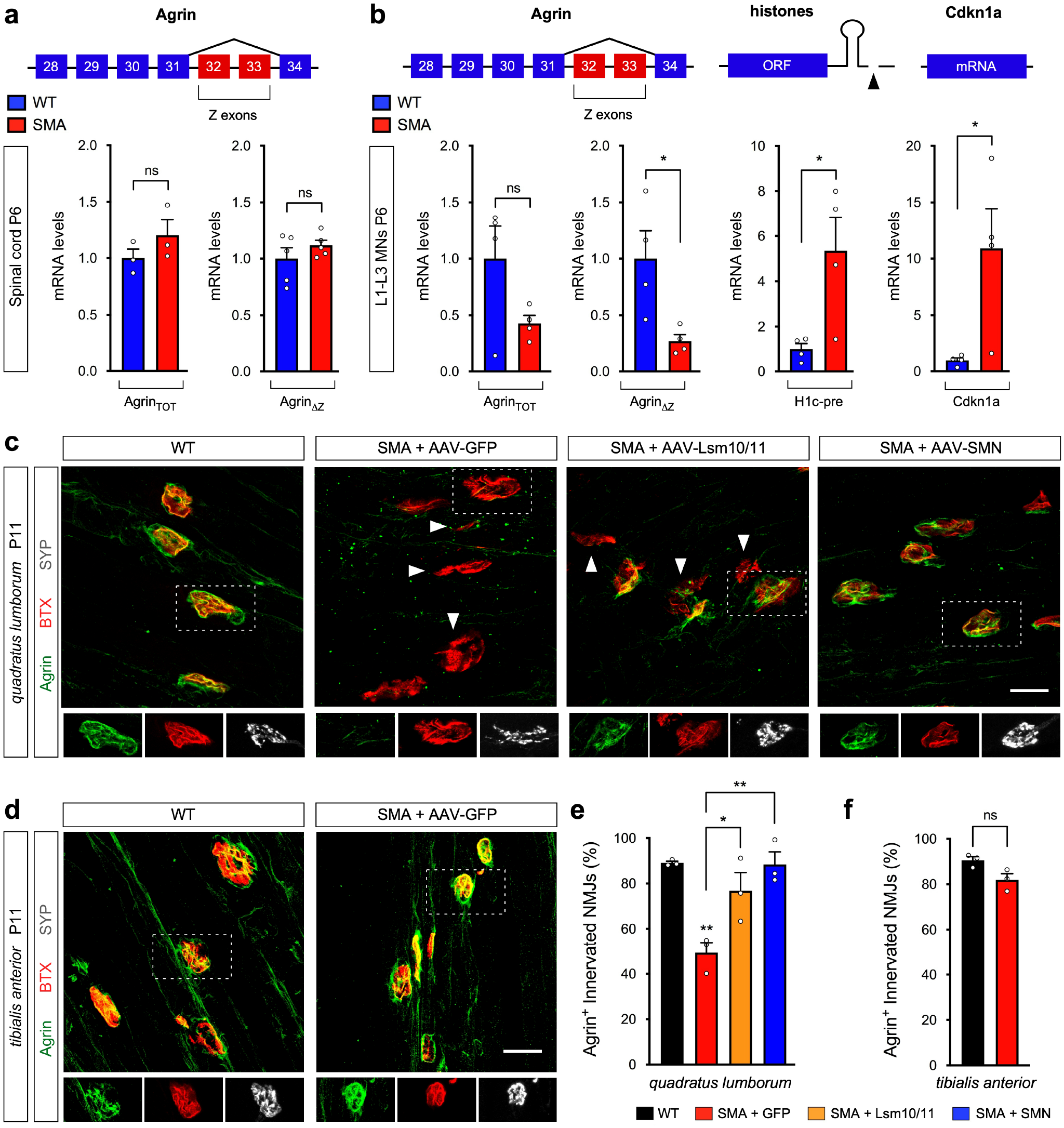
Lsm10/11 co-expression restores Agrin loss induced by SMN deficiency at vulnerable NMJs in SMA mice. **a**, RT-qPCR analysis of total (TOT) and Z exon-skipped (ΔZ) Agrin mRNAs in the spinal cord of WT and SMA mice at P11. Schematic representation of Z exons and neighboring exons of Agrin mRNA is shown. Data are mean and s.e.m (n=3 mice for Agrin_TOT_; n=5 mice for Agrin_ΔZ_) normalized to Gapdh and expressed relative to levels in WT mice set as 1. NS, not significant (P>0.05) (two-tailed Student’s *t* test; P=0.2740 for Agrin_TOT_; P=0.3055 for Agrin_ΔZ_). **b**, RT-qPCR analysis of Agrin_TOT_ and Agrin_ΔZ_ mRNAs as well as 3’-extended histone H1c pre-mRNA and Cdkn1a mRNA in laser capture microdissected L1-L3 motor neurons from WT and SMA mice at P11. Data are mean and s.e.m (n=4 independent experiments). *P<0.05 (two-tailed Student’s *t* test; P=0.0281 for Agrin_ΔZ_; P=0.0273 for H1c; P=0.0318 for Cdkn1a). NS, not significant (P>0.05) (two-tailed Student’s *t* test; P=0.1017 for Agrin_TOT_). **c**, NMJ immunostaining with Agrin, Synaptophysin (SYP) and *α*-bungarotoxin (BTX) of *quadratus lumborum* muscles from untreated WT mice and SMA mice injected with AAV9-GFP, AAV9-Lsm10/11 or AAV9-SMN at P11. SYP^+^ NMJs are indicated by arrowheads and SYP staining is only shown for one representative innervated NMJ (dotted box) in the bottom insets. Scale bar=20µm. **d**, NMJ immunostaining with Agrin, Synaptophysin (SYP) and *α*-bungarotoxin (BTX) of *tibialis anterior* muscles from the same groups as in (**c**) at P11. Denervated NMJs lacking pre-synaptic SYP staining are indicated by arrowheads and SYP staining is only shown for one representative innervated NMJ (dotted box) in the bottom insets. Scale bar=20µm. **e**, Percentage of innervated NMJs that are Agrin^+^ from experiments as in (**c**). Data are mean and s.e.m (n=3 mice). *P<0.05; **P<0.01 (one-way ANOVA with Tukey’s *post hoc* test; P=0.0036 for WT vs SMA+GFP; P=0.0287 for SMA+GFP vs SMA+Lsm10/11; P=0.0040 for SMA+GFP vs SMA+SMN). **f**, Percentage of innervated NMJs that are Agrin^+^ from experiments as in (**d**). Data are mean and s.e.m (n=3 mice). NS, not significant (P>0.05) (two-sided Student’s *t* test; P=0.0545).

We next directly investigated Agrin expression at the NMJ of vulnerable and resistant muscles from WT and SMA mice by immunostaining (Fig. 4c-f) because a reduction of Agrin at SMA NMJs has not been documented previously. Focusing on innervated NMJs labeled by both synaptophysin and *α*-bungarotoxin (BTX), we found that Agrin is absent from half of the innervated NMJs in the vulnerable QL muscle from AAV9-GFP injected SMA mice but present in nearly all NMJs from WT mice (Fig. 4c, e). Neither denervation nor loss of Agrin was observed at NMJs from the resistant tibialis anterior (TA) muscle of SMA mice (Fig. 4d, f). Thus, SMN deficiency induces selective loss of Agrin expression at NMJs that innervate vulnerable muscles in SMA mice. Importantly, Agrin expression at vulnerable SMA NMJs was restored by AAV9-Lsm10/11 in addition to AAV9-SMN (Fig. 4c, e), while the reduction in pre-synaptic SV2B levels^38^ that we also found to be specific for NMJs in the QL but not the TA of SMA mice is only restored by AAV9-SMN (Extended Data Fig. 8). Together, these results reveal that disruption of Agrin expression at the NMJ is a specific downstream consequence of U7 snRNP dysfunction induced by SMN deficiency that contributes to neuromuscular pathology in SMA mice.

## Discussion

SMN is a multifunctional protein that is essential for NMJ biology as shown by muscle denervation being a hallmark of SMA pathology in both humans and mouse models^1, 2, 8^. Through selective restoration of U7 snRNP biogenesis in SMA mice, our findings reveal a surprising role for SMN-mediated U7 snRNP assembly and histone mRNA 3’-end processing in controlling NMJ integrity through Agrin expression that is disrupted in motor neuron disease (Extended Data Fig. 9), uncovering a novel RNA-mediated pathogenic mechanism in SMA and linking U7 function to neuromuscular development. The results also establish the independent nature of the pathogenic cascades driving death and denervation of SMA motor neurons, which emerge as distinct SMN-dependent downstream events triggered by dysregulation in RNA splicing^22, 25, 26^ and histone mRNA 3’-end processing (this study), respectively. Recently, biallelic mutations in the *LSM11* and U7 (*RNU7*) genes have been linked to Aicardi–Goutières syndrome (AGS)^39^, and defective Agrin expression could contribute to some of the clinical features of AGS involving muscle dysfunction. Dysregulation of this RNA pathway could also participate in the NMJ pathology of other neurodegenerative diseases like ALS, for which there is evidence of interference with SMN function^40, 41^ and histone mRNA expression^42–44^.

Our findings provide a highly unanticipated functional connection that links seemingly disparate biological processes such as SMN-mediated U7 snRNP assembly and 3’-end processing of histone mRNAs encoding the main protein constituents of chromatin with the neurobiology of synaptic connections between motor neurons and muscle. A multitude of regulatory mechanisms contribute to establish and refine synaptic connections of motor neurons with their target muscle during development as well as sustain these connections in adulthood^45–47^. Our findings identify the pathway of SMN-dependent, U7-mediated histone mRNA processing among these mechanisms, uncovering a new facet of RNA regulation of motor neuron biology and disease. While future studies are needed to determine the precise intervening steps between histone dysregulation and reduced Agrin expression, the only cellular function of both U7 snRNP and its core subunits Lsm10 and Lsm11 is in the 3’-end cleavage of histone mRNAs^13–17^, and SMN deficiency acts cell autonomously within motor neurons to induce NMJ denervation in SMA mouse models^48, 49^. Thus, the functional requirement of U7 snRNP biogenesis and histone mRNA 3’-end processing to maintain the integrity of NMJs through Agrin highlights a previously unappreciated demand for replication-dependent histone synthesis in motor neurons that broadens the relevance of this SMN-regulated RNA pathway beyond its canonical role in the process of genome replication in dividing cells. Expanding on other facets of histone metabolism that contribute to neuronal transcription and function^50, 51^, a plausible scenario is that the pathway of SMN-mediated U7 snRNP assembly ensures an adequate supply of histones for proper chromatin function and expression of essential synaptic genes in motor neurons.

## Acknowledgements

We are grateful to Samie Jaffrey for comments and critical reading of the manuscript. We thank Brian Kaspar for providing the dsAAV-CB vector and Markus Ruegg for the kind gift of Agrin antibodies. This work was supported by the Muscular Dystrophy Association (LP) and NIH grants F31NS079002 (ST), R01NS102451 (LP), R01NS078375 (GZM.) and R01AA027079 (GZM).

## Author contributions

S.T. and L.P. conceived the study. S.T., M.V.A. and C.M.S. performed experiments. S.T., M.V.A., C.M.S., G.Z.M. and L.P. participated in experimental design and data analysis. S.T. and L.P. wrote the manuscript with contributions from all authors.

## Materials & Correspondence

Correspondence and material requests should be addressed to L.P.

## Methods

### DNA constructs

The open reading frames (ORFs) of mouse Lsm10 and Lsm11 were amplified from a cDNA library generated from NIH3T3 fibroblasts and cloned into pcDNA3 as fusions with a FLAG tag sequence at the amino terminus (Invitrogen). The resulting FLAG-tagged ORFs were then cloned into pRRL.PGK.IRES.Neo and pRRL.PGK.IRES.Hygro lentiviral vectors, which were generated by replacing the GFP sequence in the pRRL.SIN.cPPT.PGK-GFP.WPRE vector (Addgene plasmid #12252) with an IRES-Neomycin and IRES-Hygromycin cassette, respectively. To generate the CMV-driven Lsm10-2A-Lsm11 construct, DNA fragments corresponding to FLAG-Lsm10 and 2A-FLAG-Lsm11 were amplified by PCR using appropriate primers and cloned sequentially into pCDNA3. The 2A self-cleaving peptide sequence from the *Thosea asigna* virus has been previously described^52^. The entire FLAG-Lsm10-2A-FLAG-Lsm11 cassette was excised from the resulting pCDNA3 plasmid and cloned downstream of the CMV enhancer and chicken *β*-actin hybrid promoter in the dsAAV-CB plasmid harboring AAV2 inverted terminal repeats (ITRs) for the production of self-complementary AAV vectors^24^. The constructs for the expression of GFP and human SMN from the same dsAAV-CB vector were previously described^25^. All constructs were verified by DNA sequencing.

### AAV9 production

AAV9 viral vectors packaged into serotype-9 capsid were custom produced by Vector BioLabs using triple-plasmid transfection of human HEK-293 cells and purification by two rounds of CsCl gradient centrifugation. Vectors were concentrated using Amicon Ultracel centrifugal filter devices with a 30,000 nominal molecular weight limit (Millipore), titered by qPCR using the primers listed in Supplementary Table 1, quality checked by SDS/PAGE and silver staining, and stored at −80°C until use.

### Lentivirus production

Viral stocks pseudotyped with the vesicular stomatitis G protein (VSV-G) were prepared by transient co-transfection of HEK293T cells (Open Biosystems) using the ViraPower™ Lentiviral Packaging Mix (Invitrogen) following manufacturer’s instructions. Supernatant was collected 48 hours post transfection and passed through a 0.45 μm syringe filter. Lentivirus was concentrated by ultracentrifugation using a Beckman SW28 swinging bucket rotor at 19,500 rpm for 2.5 hours at 16°C, reconstituted in PBS, and stored at −80°C until use.

### Cell Lines and Treatments

The NIH3T3 (Smn_RNAi_ and SMN/Smn_RNAi_) cell lines used in this study were described previously^18, 19^. To generate stable NIH3T3 cell lines with overexpression of Lsm10 and/or Lsm11, NIH3T3-Smn_RNAi_ cells were transduced with the corresponding lentiviral vectors and selected with neomycin/G418 (0.5 mg/ml; for NIH3T3-Lsm10/Smn_RNAi_), Hygromycin B (250 μg/ml; for NIH3T3-Lsm11/Smn_RNAi_) or both antibiotics (for NIH3T3-Lsm10/Lsm11/Smn_RNAi_). For all cell lines, depletion of endogenous Smn was induced by treatment with doxycycline (100ng/ml) for 5 days prior to analysis.

### Animal procedures

All mouse work was performed in accordance with the National Institutes of Health Guidelines on the Care and Use of Animals, complied with all ethical regulations and approved by the IACUC committee of Columbia University. The SMNΔ7 mouse line FVB.Cg-Grm7^Tg(SMN2)89Ahmb^ Smn1^tm1Msd^ Tg(SMN2*delta7)4299Ahmb/J on a pure FVB/N genetic background was obtained from the Jackson Laboratory (Jax stock #005025) and used to generate SMA mice (*Smn^-/–^/SMN2^+/+^/SMNΔ7^+/+^*). Genotyping was performed using the primers listed in Supplementary Table 1 as previously described^49^. Equal proportions of mice of both sexes were used and aggregated data are presented because gender-specific differences were not found. ICV injections were performed in newborn mice anesthetized by isoflurane inhalation. A single injection was carried out using 5μl of AAV9 virus containing ∼1.0 x 10^11^ viral genomes in phosphate buffered saline (PBS) containing 5% glycerol, 0.001% Pluronic F-68, and 0.02% Fast Green dye (Sigma). After 30 min recovery, pups were placed in their breeder cage and monitored daily for survival, weight and righting time. Mice were sacrificed and tissue collection was performed in a dissection chamber under continuous oxygenation (95%O_2_/5%CO_2_) in the presence of cold (∼12°C) artificial cerebrospinal fluid (aCSF) containing 128.35mM NaCl, 4mM KCl, 0.58mM NaH_2_PO_4_, 21mM NaHCO_3_, 30mM D-Glucose, 1.5mM CaCl_2_, and 1mM MgSO_4_.

### Electrophysiology

For recordings of the monosynaptic reflex, the intact *ex vivo* spinal cord preparation was transferred to a customized recording chamber after dissection and continuously perfused with aCSF at a temperature of 21°-25°C. The dorsal root and ventral root of the L1 segment were placed into suction electrodes for stimulation or recording, respectively. The extracellular recorded potentials were recorded in response to a brief (0.2ms) orthodromic stimulation of the L1 dorsal root. The stimulus threshold was defined as the current at which the minimal evoked response was recorded in 3 out of 5 trials. Clampex (v10.2, Molecular Devices) software was used for data acquisition and Clampfit (v10.2, Molecular Devices) was used for data analysis. The monosynaptic component of the EPSP amplitude was measured from the onset of response to 3ms. Measurements were taken from averaged traces of 5 trials elicited at 0.1Hz. The function of NMJs of the QL muscle was assessed *ex vivo* as previously described^49^. After the removal of the spinal cord, the remaining vertebral column with the ventral root L1-L3 in continuity to the QL was transferred to the recording chamber containing aCSF at a temperature of 21°-25°C. The ventral root L1 was placed into a suction electrode to stimulate the motor neuron axons. Visual twitching of the QL after stimulation of the ventral root L1 ensured proper stimulation of the muscle. Subsequently, to measure the compound muscle action potential (CMAP), a concentric bipolar electrode was inserted in the QL between the insertion point of ventral roots L1 and L2. The stimulus threshold was defined as the current at which the minimal evoked response was recorded in 3 out of 5 trials. The nerve was stimulated at 1, 2, 5 and 10X threshold to ensure a supramaximal stimulation of the muscle. The maximum CMAP amplitude (baseline-to-peak) was determined as the average from 5 measurements.

### Laser capture microdissection (LCM)

Isolation of motor neurons by LCM was performed as previously described^18, 22^. Briefly, CTb conjugated to Alexa 488 was delivered by intramuscular injection in the iliopsoas and QL muscles of wild type and SMA mice at P2 using a finely pulled glass microelectrode. The L1-L3 spinal segments were dissected from the injected mice at P6, embedded in OCT, and flash frozen prior to sectioning with a cryostat. Sections (14μm) were mounted on PEN-membrane slides 2.0 (Zeiss), fixed in 100% ethanol for 15 seconds, and air dried for 30 seconds prior to microdissection of CTb^+^ motor neuron somata using a DM6000B microscope equipped with a LMD6000 laser capture unit (Leica).

### RNA analysis

Isolation of total RNA from mouse tissue and cultured cells was carried out using TRIzol reagent (Ambion) as per manufacturer’s instructions followed by treatment with RNAse-free DNaseI (Ambion). cDNA was generated from 1μg of total RNA using the RevertAid RT Reverse Transcription Kit (ThermoFisher) with a mixture of random hexamer and oligo dT primers. qPCR reactions were performed using 2.5% of the cDNA and Power SYBR Green PCR master mix (Applied Biosystem) in technical triplicates and normalized to endogenous Gapdh mRNA levels. Total RNA was purified from ∼200 motor neurons per biological replicate using the Absolutely RNA Nanoprep Kit (Agilent). Amplified cDNA was prepared from total RNA using the Ovation PicoSL WTA System V2 Kit (NuGEN) and purified with the MinElute Reaction Cleanup Kit (Qiagen). RNA quality and quantity were assessed using a 2100 Bioanalyzer (Agilent). The primers used for RT-qPCR experiments are listed in Supplementary Table 1. For Northern blot analysis, total RNA (2μg) was analyzed by electrophoresis on a denaturing 8% polyacrylamide gel containing 8 M urea followed by transfer to a positively-charge Hybond nylon membrane (GE Healthcare). Radioactive antisense RNA probes were *in vitro* transcribed from DNA oligonucleotide templates in the presence of [*α*^32^P]-UTP (3000 Ci/mmol; Perkin Elmer) using the MEGAshortscript T7 kit (Invitrogen) and purified by passing through a P-30 Tris Micro Bio-Spin Column (Bio-Rad). Prior to hybridization, labeled RNA probes were denatured at 70°C for 5 minutes and transferred to ice. Membranes were pre-hybridized for at least 30 minutes at 60°C in a rotary oven in ULTRAhyb-oligo hybridization buffer (Invitrogen). After pre-hybridization, labeled probe (1×10^6^ cpm/ml of hybridization buffer) was added to the same hybridization buffer and incubated overnight at 60°C in a rotary oven. The membrane was washed two times for 30 minutes each in 2X SSC/0.5% SDS at 60°C and then exposed to either a phosphoscreen or X-ray film. Quantification was carried out using a Typhoon PhosphorImager (Molecular Dynamics). The sequences of probes used are listed in Supplementary Table 1.

### Protein analysis

Total protein extracts were generated by cell lysis in SDS sample buffer (2% SDS, 10% glycerol, 5% *β*-mercaptoethanol, 60mM Tris-HCl pH 6.8, and bromophenol blue), followed by brief sonication and boiling. Protein was quantified using the *RC DC*^TM^ Protein Assay (Bio-Rad). For immunoprecipitation experiments, cell extracts were prepared by homogenization in ice cold RSB-100 buffer (100mM NaCl, 10mM Tris-HCl pH 7.4, 2.5mM MgCl_2_) containing 0.1% NP40, protease inhibitors (Pierce/Thermo Scientific), and phosphatase inhibitors (PhosSTOP; Roche/Sigma-Aldrich), followed by passing five times through a 27-gauge needle and centrifugation at 10,000 x g for 15 minutes at 4°C. Extract supernatant was quantified using the Bradford Quick Dye assay (Bio-Rad). Antibodies were bound to protein G-Sepharose (Sigma-Aldrich) in RSB-100 buffer containing 0.1% NP40, protease inhibitors, and phosphatase inhibitors for 2 hours at 4°C. Following five washes with the same buffer, antibody-bound beads were incubated with 200 μg of cell extract for 2 hours at 4°C with tumbling. Following five washes with the same buffer, samples were processed for either RNA or protein analysis. Bound RNAs were extracted by treatment with proteinase K (200 μg) diluted in proteinase K buffer (10 mM EDTA, 100 mM Tris-HCl pH 7.5, 300 mM NaCl, 2% SDS) for 20 minutes at room temperature with gentle agitation followed by phenol-chloroform extraction and ethanol precipitation. Immunoprecipitated RNA was analyzed by electrophoresis on a denaturing 8% polyacrylamide gel containing 8 M urea followed by Northern blotting. Proteins were eluted from beads by boiling in SDS sample buffer and analyzed by SDS/PAGE on 12% polyacrylamide gels followed by Western blotting as previously described^19^.

### *In vitro* snRNP assembly

Radioactive U1 and U7 snRNAs were generated by run-off transcription with T7 polymerase from linearized plasmid DNA templates in the presence of m^7^G cap analog (Promega) using the MEGAshortscript T7 kit (Invitrogen) as previously described^14, 53, 54^. Cell and tissue extracts were prepared in ice-cold reconstitution buffer (20mM HEPES-KOH, pH 7.9, 50mM KCl, 5mM MgCl_2_, 0.2mM EDTA, 5% glycerol) containing 0.01% NP-40 and *in vitro* snRNP assembly reactions were carried out for 1 hour at 30°C as previously described^14, 53, 54^. Following treatment with heparin sulfate (5mg/ml) and 2M urea for 15 minutes at room temperature, reactions were immunoprecipitated with anti-SmB (18F6) antibodies for 2 hours at 4°C in RSB-500 buffer (500mM NaCl, 10mM Tris-HCl, pH 7.4, 2.5mM MgCl_2_) containing 0.1% NP40, protease inhibitors (Pierce/Thermo Scientific), and phosphatase inhibitors (PhosSTOP; Roche/Sigma-Aldrich). Following antibody conjugation, beads were washed 5 times in RSB-500 buffer. Immunoprecipitated snRNAs waere analyzed by electrophoresis on a denaturing 8% polyacrylamide gel containing 8 M urea followed by autoradiography. Quantification was done using a Typhoon PhosphorImager (Molecular Dynamics).

### Immunohistochemistry and confocal microscopy

Freshly dissected spinal cords were immersion-fixed in 4% paraformaldehyde (PFA) in PBS overnight at 4°C. Specific lumbar segments were identified by the ventral roots, embedded in 5% agar and sectioned at 75μm with a VT1000S Vibratome (Leica). Sections were washed briefly in PBS and then blocked for 2 hours at room temperature in 10% normal donkey serum (Millipore) in 0.01M PBS containing 0.4% Triton X-100, pH 7.4. Immunostaining, washing, and mounting were then performed as previously described^21^. For NMJ analysis, freshly dissected QL and TA muscles were immersion-fixed in 4% PFA-PBS for 1 hour at room temperature, followed by a PBS wash and cryoprotection in 30% sucrose overnight at 4°C. Muscle was then embedded in Optimal Cutting Temperature (OCT) compound (Fisher), frozen on dry ice, and sectioned at 30μm with a CM3050S cryostat (Leica) for processing by immunostaining as previously described (Van Alstyne 2021). The antibodies used for these experiments are listed in Supplementary Table 2. For myofiber analysis, freshly dissected triceps muscle was immersion-fixed in 4% PFA-PBS for 1 hour at room temperature followed by a PBS wash. Muscle was then cryoprotected by incubation with a gradient of sucrose solutions from 10% sucrose to 30% sucrose in 0.1M phosphate buffer, pH 7.4 for 1 hour each time. Cryoprotected tissue was then removed from sucrose, embedded in OCT, and flash-frozen in liquid nitrogen-cooled 2-methyl butane (Sigma) prior to serial sectioning at 25μm thickness onto Superfrost Plus glass slides (Fisher) using a CM3050S cryostat (Leica). Slides were incubated in TBS containing 0.2% Triton-X for 30 minutes and then with a solution containing TRITC-conjugated phalloidin (Sigma-Aldrich) and DAPI (ThermoFisher Scientific) overnight at 4°C. The following day, slides were washed 3 times with TBS-T and mounted with a glass coverslip using Fluoromount-G (Southern Biotech) mounting media. All images were collected with an SP5 confocal microscope (Leica) running the LAS AF software (v2.5.2.6939) and analyzed off-line using the Leica LAS X software (v1.9.0.13747) from z-stack images. For motor neuron number quantification, images were acquired from all the 75μm sections of L1 and L5 spinal segments using a 40X objective at 3μm steps in the z-axis steps. Images for H2AX intensity analysis were acquired using a 40X objective at identical settings for WT and SMA samples at 3μm steps and quantified by measuring the peak intensity per unit area of the nuclear region of motor neurons as previously described^21, 22^. For VGluT1^+^ synapses quantification, images were acquired using a 40X objective throughout the entire thickness of 75μm L1 sections at 0.2μm z-steps. The total number of VGluT1^+^ synapses on soma was determined by counting all the corresponding inputs on the surface of each motor neuron cell body. Images for NMJ analysis were acquired using a 20X objective at 2μm steps in the z-axis. Myofiber area was quantified using Fiji (v1.0) and manual identification of myofiber edges.

### Statistical analysis

Differences between two groups were analyzed by two-tailed unpaired Student’s *t*-test and differences among three or more groups were analyzed by one-way ANOVA followed by the Tukey’s *post hoc* tests for multiple comparisons as indicated. GraphPad Prism (v9.1.2) was used for all statistical analyses and P values are indicated as follows: *P<0.05; **P<0.01; ***P<0.001; **** P< 0.0001. Exact P values are listed in the figure legends.

---

**Extended Data Fig. 1.**
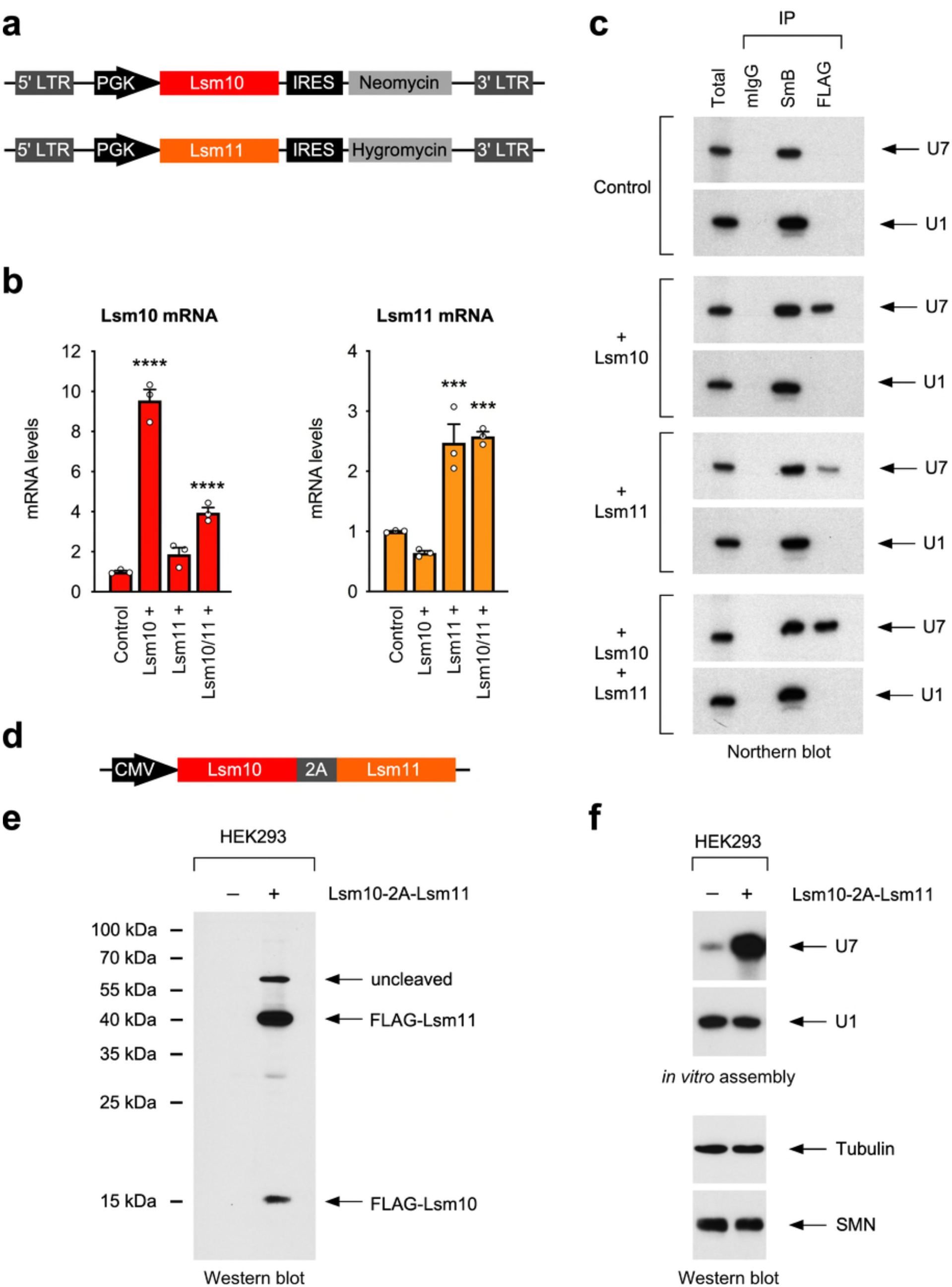
Co-expression of Lsm10 and Lsm11 enhances U7 snRNP assembly. **a**, Schematic representation of the lentiviral constructs used to generate stable NIH3T3 cell lines with mouse Lsm10 and/or Lsm11 overexpression. **b**, RT-qPCR analysis of Lsm10 and Lsm11 mRNA levels in NIH3T3-Smn_RNAi_ cells with and without (control) overexpression of Lsm10 and Lsm11 either individually or in combination. Data are mean and s.e.m (n=3 independent experiments) normalized to Gapdh mRNA and expressed relative to control NIH3T3-Smn_RNAi_ cells set as 1. ***P<0.001, ****P<0.0001 (one-way ANOVA with Tukey’s *post hoc* test; Lsm10 mRNA analysis: P<0.0001 for Lsm10; P<0.0001 for Lsm10/11; Lsm11 mRNA analysis: P=0.0008 for Lsm11; P=0.0005 for Lsm10/11). **c**, Northern blot analysis of U1 and U7 snRNAs immunoprecipitated with the indicated antibodies from extracts of NIH3T3-Smn_RNAi_ cell lines with and without overexpression of Lsm10 and Lsm11 either individually or in combination. **d**, Schematic representation of the vector expressing both Lsm10 and Lsm11 with an intervening 2A self-cleaving peptide. **e**, Western blot analysis with anti-FLAG antibodies of HEK293T cells with or without transient transfection of the Lsm10-2A-Lsm11 vector depicted in (**d**). **f**, *In vitro* snRNP assembly for U7 and U1 snRNAs (top panels) and Western blot analysis of SMN levels (bottom panels) using the same extracts as in (**e**).

**Extended Data Fig. 2.**
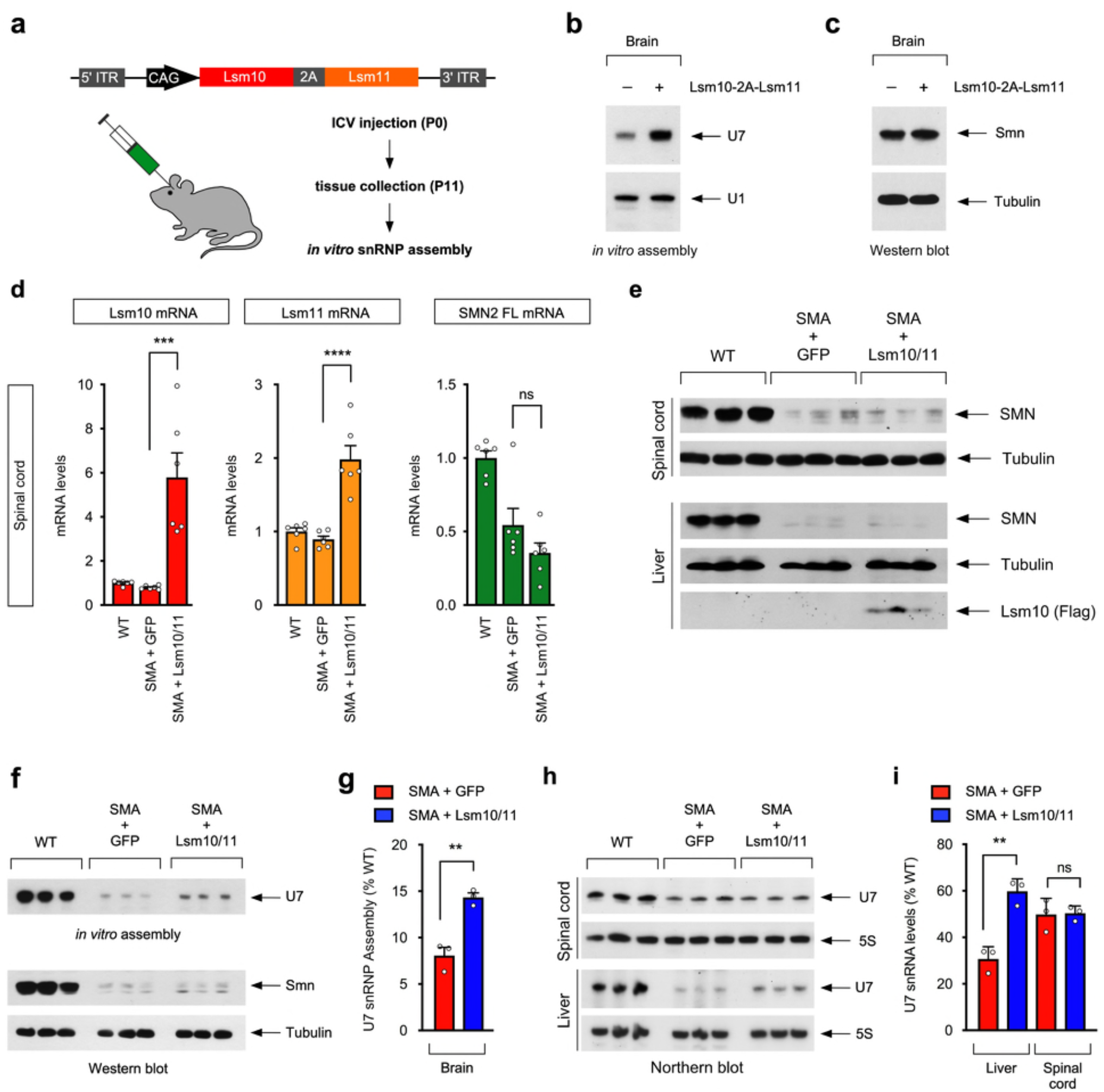
Co-expression of Lsm10 and Lsm11 does not increase SMN expression and enhances U7 snRNP biogenesis in SMA mice. **a**, Schematic representation of the AAV9 viral vector and workflow to achieve Lsm10 and Lsm11 co-expression *in vivo*. **b**, *In vitro* snRNP assembly of U1 and U7 snRNAs in brain extracts from uninjected and AAV9-Lsm10-2A-Lsm11 injected WT mice at P11. **c**, Western blot analysis of SMN levels in brain extracts from the same samples used in (**b**). **d**, RT-qPCR analysis of Lsm10 (left), Lsm11 (middle), and full-length human SMN2 (right) mRNA levels in the spinal cord of uninjected WT mice and SMA mice injected with AAV9-GFP or AAV9-Lsm10/11 at P11. Data are mean and s.e.m (n=6 mice) normalized to Gapdh mRNA and expressed relative to WT set as 1. ***P<0.001, ****P<0.0001 (one-way ANOVA with Tukey’s *post hoc* test; Lsm10 mRNA analysis: P=0.0002 for SMA+GFP vs SMA+Lsm10/11; Lsm11 mRNA analysis: P<0.0001 for SMA+GFP vs SMA+Lsm10/11). NS, not significant (P>0.05) (one-way ANOVA with Tukey’s *post hoc* test; SMN2 FL analysis: P=0.2487 for SMA+GFP vs SMA+Lsm10/11). **e**, Western blot analysis of SMN levels in spinal cord and liver from the same groups as in (**d**). Flag-Lsm10 could only be detected with anti-FLAG antibodies from liver but not spinal cord of SMA mice injected with AAV9-Lsm10/11. **f**, *In vitro* U7 snRNP assembly (top panel) and Western blot analysis (bottom panels) of brain extracts from uninjected WT mice and SMA mice injected with AAV9-GFP or AAV9-Lsm10/11 at P11. **g**, Percentage of U7 snRNP assembly relative to uninjected WT mice from the experiment in (**f**). Data are mean and s.e.m (n=3 mice). **P<0.01 (two-tailed Student’s *t* test; P=0.0033). **h**, Northern blot analysis of U7 snRNA and 5S rRNA in spinal cord (top panels) and liver (bottom panels) from the same groups as in (**f**). **i**, Percentage of U7 snRNA levels relative to uninjected WT mice from the experiment in (**h**). Data are mean and s.e.m (n=3 mice). **P<0.01 (two-tailed Student’s *t* test; P=0.0027). NS, not significant (P>0.05) (two-tailed Student’s *t* test; P=0.9060).

**Extended Data Fig. 3.**
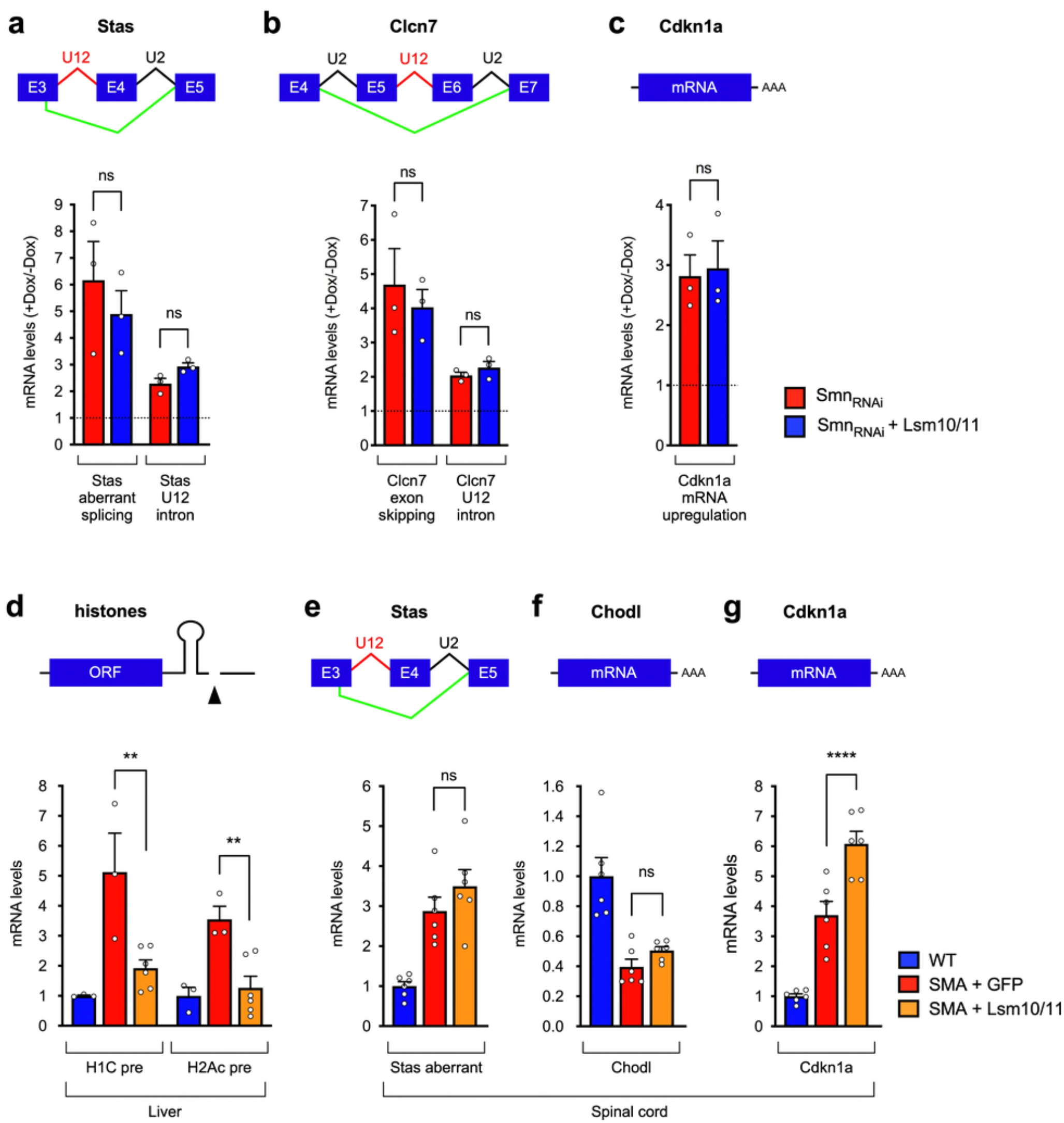
Co-expression of Lsm10 and Lsm11 does not correct U7-independent RNA processing events induced by SMN deficiency. **a**, RT-qPCR analysis of aberrant U12 splicing and U12 intron retention in the Stasimon (Stas) mRNA induced by SMN deficiency in NIH3T3-Smn_RNAi_ and NIH3T3-Lsm10/11/Smn_RNAi_ cells. Schematic representation of the monitored RNA processing events is shown at the top. For each cell line, fold change values in Dox treated cells relative to untreated cells set as 1 are shown. Data are mean and s.e.m (n=3 independent experiments) normalized to Gapdh mRNA. NS, not significant (P>0.05) (two-tailed Student’s *t* test; P=0.4952 for aberrant splicing; P=0.0547 for U12 intron retention). **b**, RT-qPCR analysis of exon skipping and U12 intron retention in the Clcn7 mRNA in the same cell lines as in (**a**). Schematic representation of the monitored RNA processing events is shown at the top. Data are mean and s.e.m (n=3 independent experiments) normalized to Gapdh mRNA. NS, not significant (P>0.05) (two-tailed Student’s *t* test; P=0.6035 for exon skipping; P=0.3143 for U12 intron retention). **c**, RT-qPCR analysis of Cdkn1a mRNA levels in the same cell lines as in (**a**). Data are mean and s.e.m (n=3 independent experiments) normalized to Gapdh mRNA. NS, not significant (P>0.05) (two-tailed Student’s *t* test; P=0.8309). **d**, RT-qPCR analysis of 3’-extended histone mRNAs in liver from uninjected WT mice and SMA mice injected with AAV9-GFP or AAV9-Lsm10/11 at P11. Schematic representation of a histone pre-mRNA and the 3’-end cleavage is shown at the top. Data are mean and s.e.m (n=3 mice for WT and SMA+GFP; n=6 mice for SMA+Lsm10/11) normalized to Gapdh mRNA. **P<0.01 (one-way ANOVA with Tukey’s *post hoc* test; P=0.0097 for H1c; P=0.0087 for H2Ac). **e-g**, RT-qPCR analysis of aberrant U12 splicing of Stas mRNA (**e**) as well as Chodl (**f**) and Cdkn1a (**g**) total mRNA levels in spinal cord from the same groups as in (**d**). Schematic representation of the monitored RNA processing events is shown at the top. Data are mean and s.e.m (n=6 mice) normalized to Gapdh mRNA. ****P<0.0001 (one-way ANOVA with Tukey’s *post hoc* test; P<0.0001 for Cdkn1a). NS, not significant (P>0.05) (one-way ANOVA with Tukey’s *post hoc* test; P=0.3819 for Stas aberrant splicing; P=0.6058 for Chodl).

**Extended Data Fig. 4.**
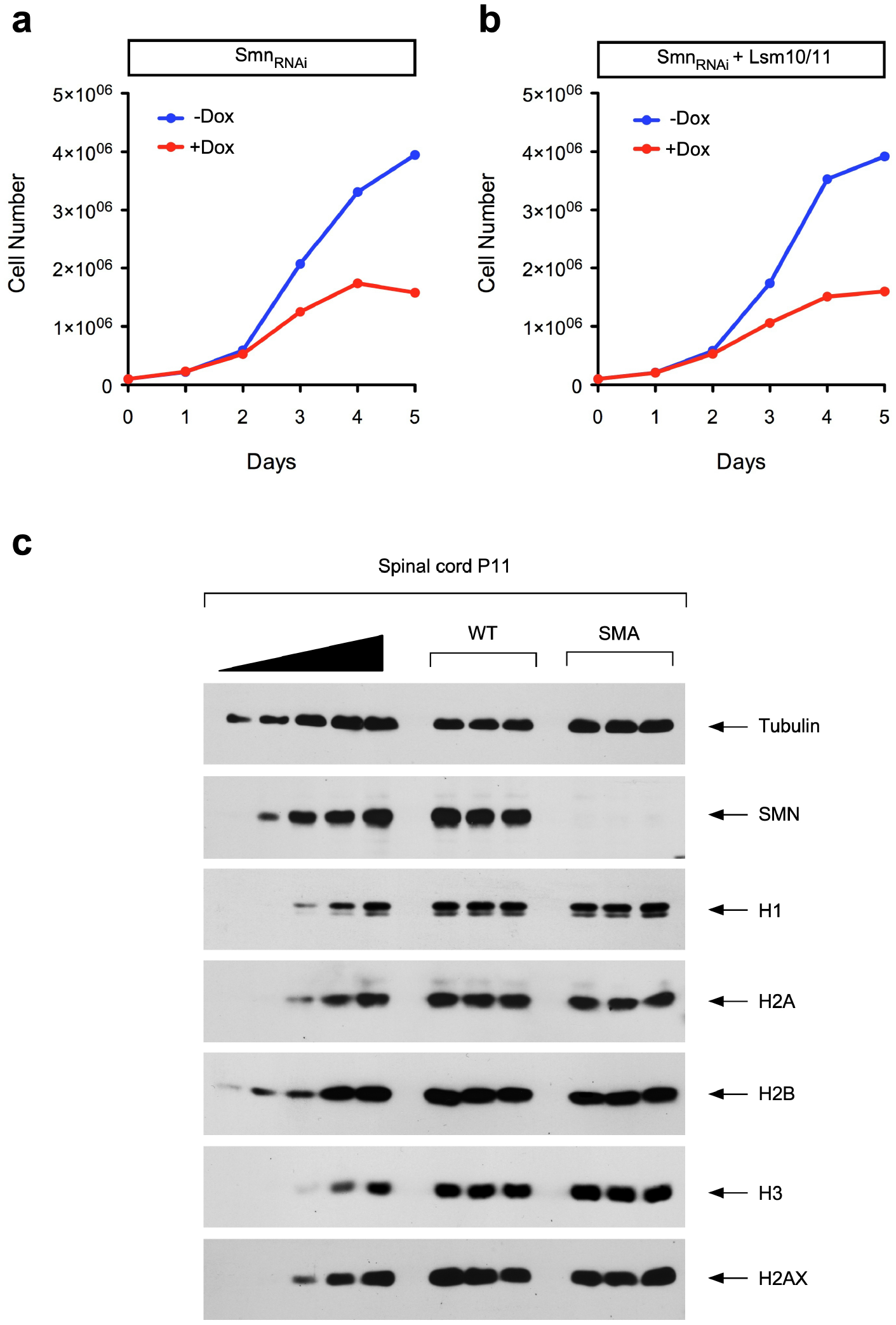
Co-expression of Lsm10 and Lsm11 does not correct cell proliferation defects in SMN-deficient mammalian cells. **a-b**, Analysis of cell proliferation in NIH3T3-Smn_RNAi_ (**a**) and NIH3T3-Lsm10/11/Smn_RNAi_ (**b**) cell lines. Equal numbers of each cell line were cultured with or without Dox for the indicated number of days and cell number was determined at each time point. **c**, Western blot analysis of histone protein levels in spinal cord from WT and SMA mice at P11. A two-fold serial dilution of WT extract is shown on the left.

**Extended Data Fig. 5.**
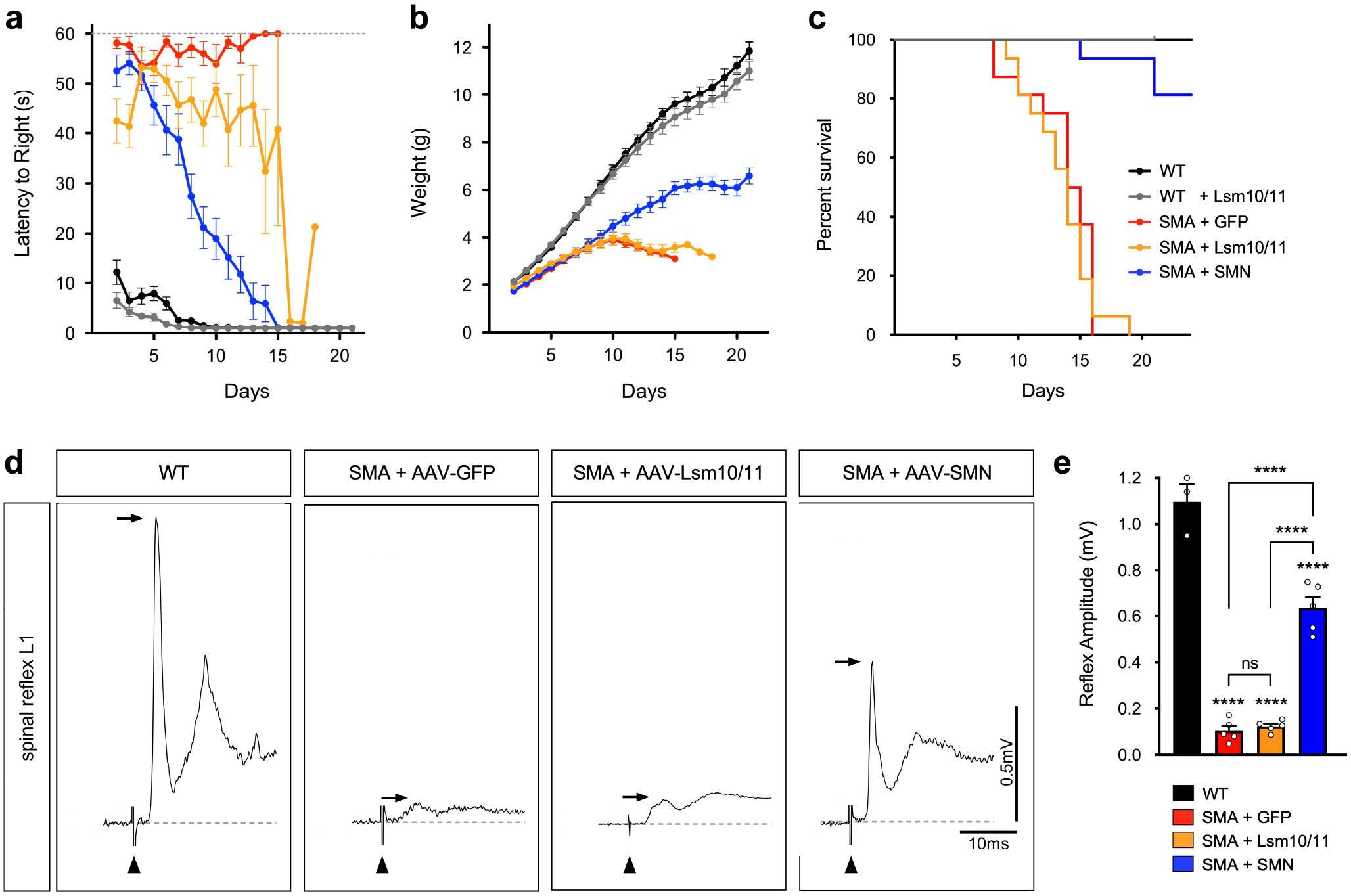
Co-expression of Lsm10 and Lsm11 enhances motor function in SMA mice. **a-b**, Righting time (**a**) and body weight (**b**) of WT mice either uninjected or injected with AAV9-Lsm10/11 and SMA mice injected with AAV9-GFP, AAV9-Lsm10/11 or AAV9-SMN. Data are mean and s.e.m from the following number of mice per group (n=16 for WT, SMA+GFP, SMA+SMN, and SMA+Lsm10/11; n=10 for WT+Lsm10/11). **c**, Kaplan-Meier plot of mouse survival from the same treatment groups and number of mice as in (**a-b**). **d**, Representative traces of spinal reflexes recorded from the L1 ventral root following L1 dorsal root stimulation in uninjected WT mice and SMA mice injected with AAV9-GFP, AAV9-Lsm10/11 or AAV9-SMN at P11. Arrows indicate peak amplitude and arrowheads indicate the stimulus artifact. **e**, Spinal reflex amplitude from experiments as in (**d**). Data are mean and s.e.m (n=3 mice for WT; n=5 for each SMA injected group). ***P<0.001 (one-way ANOVA with Tukey’s *post hoc* test; P<0.0001 for WT vs SMA+GFP; P<0.0001 for WT vs SMA+Lsm10/11; P<0.0001 for WT vs SMA+SMN; P<0.0001 for SMA+GFP vs SMA+SMN; P<0.0001 for SMA=Lsm10/11 vs SMA+SMN). NS, not significant (P>0.05) (one-way ANOVA with Tukey’s *post hoc* test; P=0.9804 for SMA+GFP vs SMA+Lsm10/11).

**Extended Data Fig. 6.**
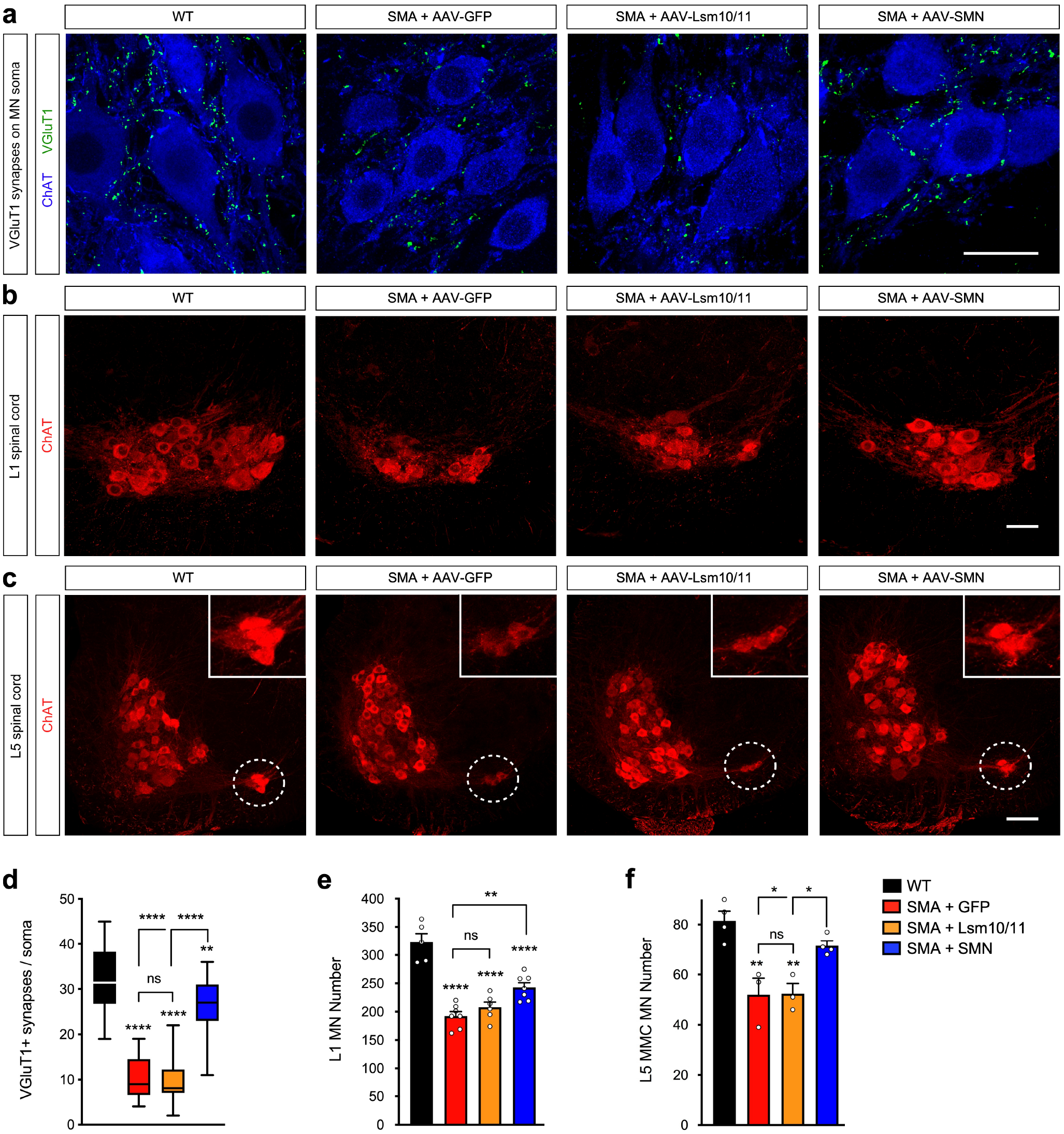
Co-expression of Lsm10 and Lsm11 does not correct the loss of proprioceptive synapses and motor neuron death in SMA mice. **a**, VGluT1 and ChAT immunostaining of L1 spinal cords from uninjected WT mice and SMA mice injected with AAV9-GFP, AAV9-Lsm10/11 or AAV9-SMN at P11. Scale bar=20µm. **b**, ChAT immunostaining of L1 spinal cords from the same groups as in (**a**). Scale bar=50µm. **c**, ChAT immunostaining of L5 spinal cords from the same groups as in (**a**). L5 medial motor column (MMC) motor neurons are indicated by the dashed circle and shown magnified in the inset in (**c**). Scale bar=125µm. **d**, Number of VGluT1^+^ synapses on the somata of L1 motor neurons from experiments as in (**a**). The box-and-whiskers graph shows the median, interquartile range, minimum and maximum from the following number of neurons and mice per group (n=22 neurons, n=3 animals for WT; n=25 neurons, n=3 animals for SMA+GFP; n=26 neurons, n=3 animals for SMA+Lsm10/11; n=21 neurons, n=3 animals for SMA+SMN). **P<0.01; ****P<0.0001 (one-way ANOVA with Tukey’s *post hoc* test; P<0.0001 for WT vs SMA+GFP; P<0.0001 for WT vs SMA+Lsm10/11; P=0.006 for WT vs SMA+SMN; P<0.0001 for SMA+GFP vs SMA+SMN; P<0.0001 for SMA+Lsm10/11 vs SMA+SMN). NS, not significant (P>0.05) (one-way ANOVA with Tukey’s *post hoc* test; P=0.9844 for SMA+GFP vs SMA+Lsm10/11). **e**, Total number of L1 motor neurons from experiments as in (**b**). Data are mean and s.e.m (n=5 mice for WT; n=7 SMA+GFP, and SMA+SMN; n=6 for SMA+Lsm10/11). **P<0.01; ****P<0.0001 (one-way ANOVA with Tukey’s *post hoc* test; P<0.0001 for WT vs SMA+GFP; P<0.0001 for WT vs SMA+Lsm10/11; P<0.0001 for WT vs SMA+SMN; P=0.0048 for SMA+GFP vs SMA+SMN). NS, not significant (P>0.05) (one-way ANOVA with Tukey’s *post hoc* test; P=0.6660 for SMA+GFP vs SMA+Lsm10/11). **f**, Total number of L5 MMC motor neurons from experiments as in (**c**). Data are mean and s.e.m (n=4 mice for WT, and SMA+SMN; n=3 for SMA+GFP, and SMA+Lsm10/11). *P<0.05; **P<0.01 (one-way ANOVA with Tukey’s *post hoc* test; P=0.0023 for WT vs SMA+GFP; P=0.0025 for WT vs SMA+Lsm10/11; P=0.032 for SMA+GFP vs SMA+SMN; P=0.0351 for SMA+Lsm10/11 vs SMA+SMN). NS, not significant (P>0.05) (one-way ANOVA with Tukey’s *post hoc* test; P>0.9999 for SMA+GFP vs SMA+Lsm10/11).

**Extended Data Fig. 7.**
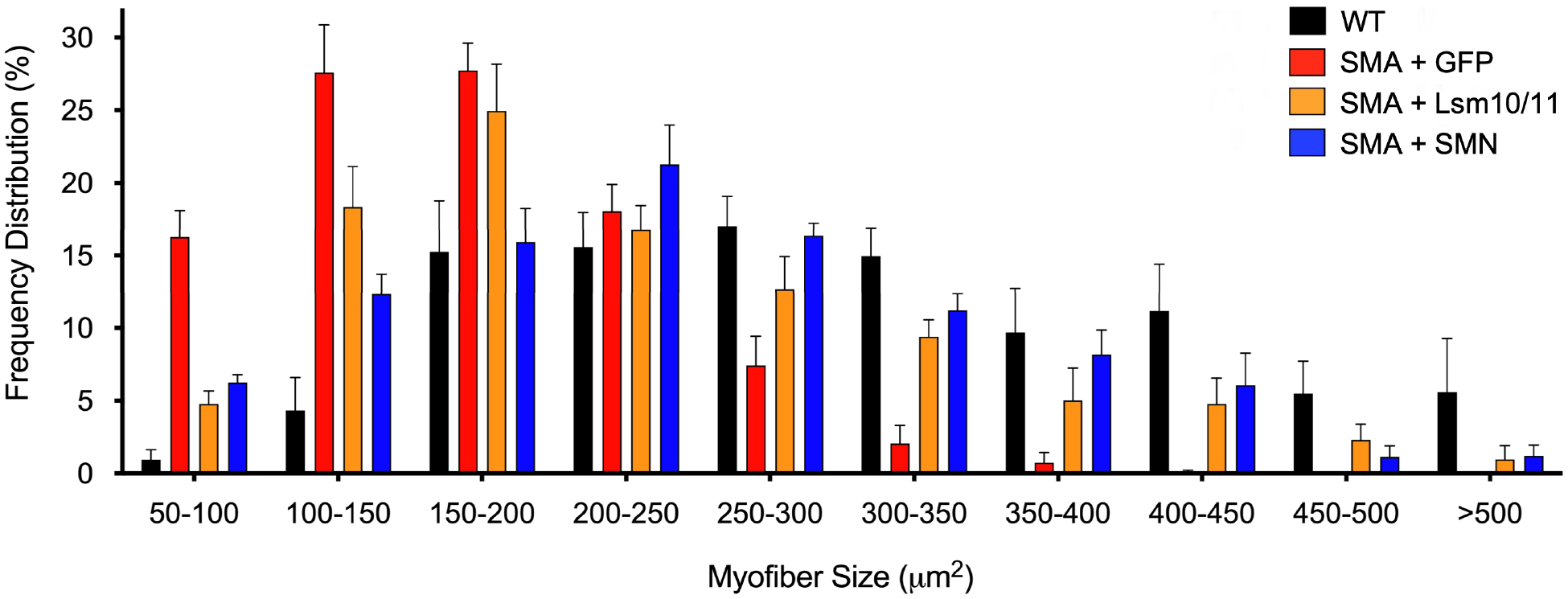
Co-expression of Lsm10 and Lsm11 improves skeletal muscle atrophy in SMA mice. Frequency distribution of myofiber sizes in the triceps muscle from uninjected WT mice and SMA mice injected with AAV9-GFP, AAV9-Lsm10/11 or AAV9-SMN at P11. Data are mean and s.e.m (n=5 mice for WT, SMA+Lsm10/11, and SMA+SMN; n=6 SMA+GFP).

**Extended Data Fig. 8.**
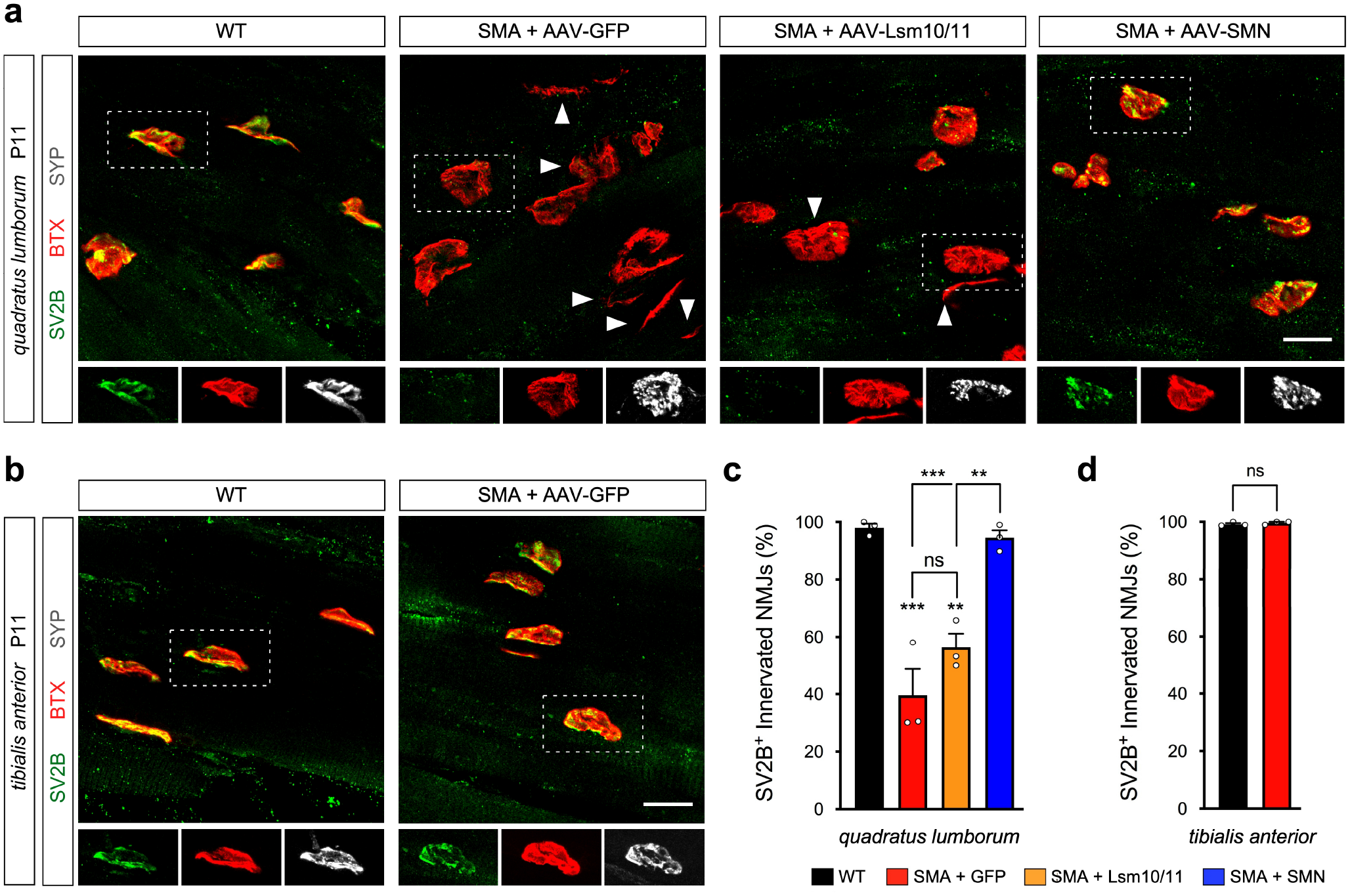
Loss of SV2B expression at vulnerable NMJs in SMA mice is not U7-dependent. **a**, NMJ immunostaining with SV2B, Synaptophysin (SYP) and *α*-bungarotoxin (BTX) of *quadratus lumborum* muscles from untreated WT mice and SMA mice injected with AAV9-GFP, AAV9-Lsm10/11 or AAV9-SMN at P11. SYP^+^ NMJs are indicated by arrowheads and SYP staining is only shown for one representative innervated NMJ (dotted box) in the bottom insets. Scale bar=20µm. **b**, NMJ immunostaining with SV2B, Synaptophysin (SYP) and *α*-bungarotoxin (BTX) of *tibialis anterior* muscles from untreated WT mice and SMA mice injected with AAV9-GFP at P11. Denervated NMJs lacking pre-synaptic SYP staining are indicated by arrowheads and SYP staining is only shown for one representative innervated NMJ (dotted box) in the bottom insets. Scale bar=20µm. **c**, Percentage of innervated NMJs that are SV2B^+^ from experiments as in (**a**). Data are mean and s.e.m (n=3 mice). **P<0.01; ***P<0.001 (one-way ANOVA with Tukey’s *post hoc* test; P=0.0003 for WT vs SMA+GFP; P=0.0028 for WT vs SMA+Lsm10/11; P=0.0004 for SMA+GFP vs SMA+SMN; P=0.0048 for SMA+Lsm10/11 vs SMA+SMN). NS, not significant (P>0.05) (one-way ANOVA with Tukey’s *post hoc* test; P=0.2071 for SMA+GFP vs SMA+Lsm10/11). **d**, Percentage of innervated NMJs that are SV2B^+^ from experiments as in (**b**). Data are mean and s.e.m (n=3 mice). NS, not significant (P>0.05) (two-sided Student’s *t* test; P=0.4267).

**Extended Data Fig. 9.**
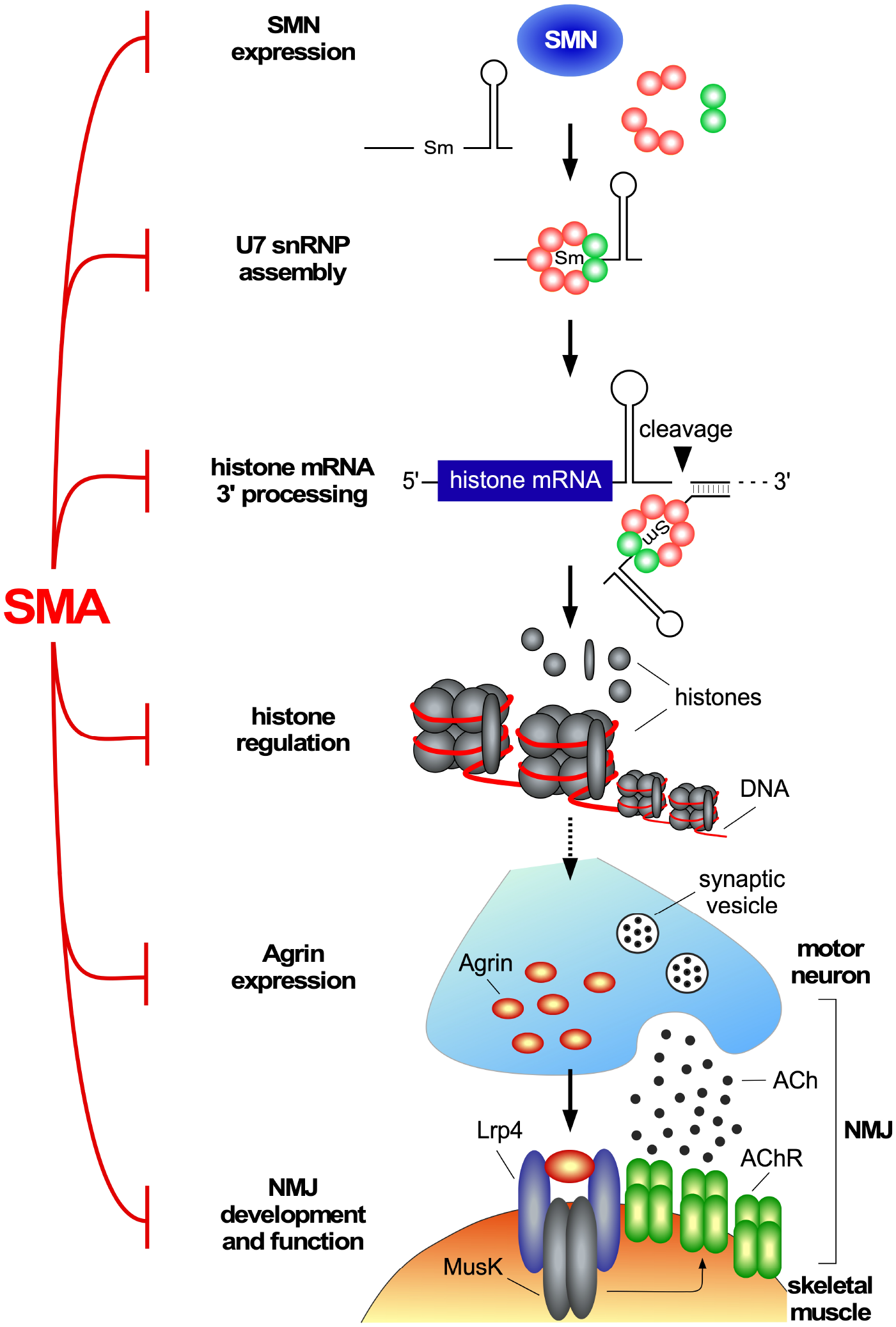
A model illustrating how U7 snRNP biogenesis dysfunction induced by SMN deficiency disrupts NMJ innervation in SMA. SMN controls normal assembly and biogenesis of U7 snRNP, which is required for proper 3’-end processing of histone mRNAs. In SMA, SMN deficiency disrupts U7 snRNP biogenesis leading to downstream dysregulation of histone gene expression including within post-mitotic motor neurons. Through mechanisms that remain to be established (dotted arrow), altered histone expression affects the normal expression of Agrin within motor neurons leading to reduced Agrin release at vulnerable NMJs, which in turn contributes to denervation and neuromuscular pathology in SMA mice.

**Supplementary Table 1.**
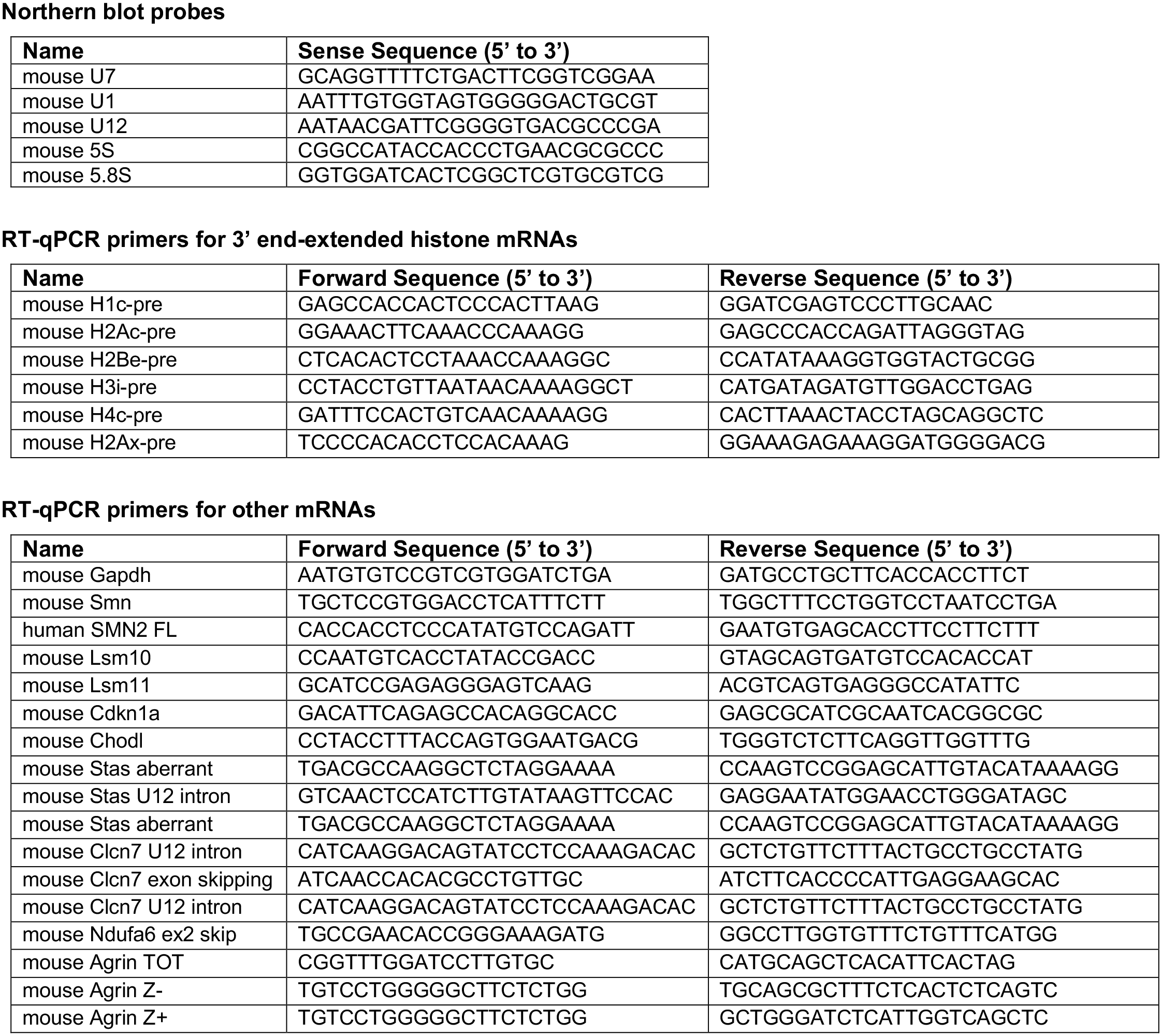
List of primers and probes used in this study.

**Supplementary Table 2.** List of antibodies used in this study.

| **Name** | **Source** | **Cat #** | **Host** | **Application** | **Dilution** |
| --- | --- | --- | --- | --- | --- |
| SMN (clone 8) | BD Transduction Lab | 610646 | Mouse | WB | 1:10,000 |
| Tubulin (DM1A) | Sigma | T9026 | Mouse | WB | 1:10,000 |
| H2A | Millipore | 07-146 | Rabbit | WB | 1:2,000 |
| H2AX | Cell Signaling | 7631S | Rabbit | WB / IHC | 1:1,000 / 1:200 |
| H2A.Z | Cell Signaling | 2718S | Rabbit | WB | 1:1,000 |
| H2B | Abcam | ab1790 | Rabbit | WB | 1:40,000 |
| H3 | Abcam | ab1791 | Rabbit | WB | 1:40,000 |
| H4 | Millipore | 07-108 | Rabbit | WB | 1:1,000 |
| Histones (pan) | Millipore | MAB052 | Mouse | WB | 1:10,000 |
| SmB (18F6) | Custom made | N/A | Mouse | IP | N/A |
| FLAG M2 | Sigma | F3165 | Mouse | WB / IP | 1:2,000 |
| ChAT | Millipore | AB144P | Goat | IHC | 1:100 |
| Neurofilament M | Millipore | AB1987 | Rabbit | IHC | 1:500 |
| VGluT1 | Custom made | N/A | Guinea pig | IHC | 1:5,000 |
| Synaptophysin | Synaptic Systems | 101-004 | Guinea pig | IHC | 1:500 |
| GFP | Sigma | G1544 | Rabbit | WB | 1:5,000 |
| GFP | Aves | GFP-1020 | Chicken | IHC | 1:500 |
| Agrin | Custom made | N/A | Rabbit | IHC | 1:1,000 |
| SV2B | Synaptic Systems | 119-102 | Rabbit | IHC | 1:200 |
| Bungarotoxin | Invitrogen | T1175 | Chicken | WB | 1:5,000 |
| Mouse immunoglobulin (IgG) | Sigma | I8765 | Mouse | IP | N/A |
| Alexa-Fluor 488 donkey anti-rabbit | Jackson | 711-545-152 | Donkey | IHC | 1:250 |
| Alexa-Fluor 488 donkey anti-goat | Jackson | 705-545-147 | Donkey | IHC | 1:250 |
| Alexa-Fluor 488 donkey anti-chicken | Jackson | 703-545-155 | Donkey | IHC | 1:250 |
| Cy3 donkey anti-rabbit | Jackson | 711-165-152 | Donkey | IHC | 1:250 |
| Cy3 donkey anti-mouse | Jackson | 715-165-150 | Donkey | IHC | 1:250 |
| Cy3 donkey anti-goat | Jackson | 705-165-147 | Donkey | IHC | 1:250 |
| Cy5 donkey anti-goat | Jackson | 705-175-147 | Donkey | IHC | 1:250 |
| Cy5 donkey anti-guinea pig | Jackson | 706-175-148 | Donkey | IHC | 1:250 |
| HRP goat anti-mouse | Jackson | 115-035-044 | Goat | WB | 1:10,000 |
| HRP goat anti-rabbit | Jackson | 111-035-003 | Goat | WB | 1:10,000 |

